# T-cell receptors identified by a personalized antigen-agnostic screening approach target shared neoantigen KRAS Q61H

**DOI:** 10.1101/2024.09.19.612910

**Authors:** Volker Lennerz, Christoph Doppler, Martina Fatho, Anja Dröge, Sigrid Schaper, Kristin Gennermann, Nadine Genzel, Stephanie Plassmann, David Weismann, Samuel W. Lukowski, Dominik Bents, Christina Beushausen, Karen Kriese, Hermann Herbst, Volkhard Seitz, Rudolf Hammer, Paul J. Adam, Stephan Eggeling, Catherine Wölfel, Thomas Wölfel, Steffen Hennig

## Abstract

Adoptive cell therapy (ACT) with TCR-engineered T-cells represents a promising alternative to TIL- or CAR-T therapies for patients with advanced solid cancers. Currently, selection of therapeutic TCRs critically depends on knowing the target antigens, a condition excluding most patients from treatment. Direct antigen-agnostic identification of tumor-specific T-cell clonotypes and TCR-T manufacturing using their TCRs can advance ACT for patients with aggressive solid cancers. We present a method to identify tumor-specific clonotypes from surgical specimens by comparing TCRβ-chain repertoires of TILs and adjacent tissue-resident lymphocytes. In seven NSCLC-patients, tumor-specific clonotypes were selected based on TIL-abundance and high tumor-to-nontumor frequency ratios. In two of the patients, we demonstrate that predicted tumor-specific clonotypes reacted against autologous tumors. In a third patient, we engineered TCR T-cells with four candidate tumor-specific TCRs that showed reactivity against the patient’s tumor and HLA-matched NSCLC cell lines. The TCR-T cells were then used to screen for candidate neoantigens and aberrantly expressed antigens. Three TCRs recognized recurrent driver-mutation KRAS Q61H-peptide ILDTAG**<u>H</u>**EEY presented by HLA-A*01:01. The TCRs were also dominant in a tumor relapse, one was found in cell free DNA. The finding of homologous TCRs in independent KRAS Q61H-positive cancers suggests a therapeutic opportunity for HLA-matched patients with KRAS Q61H-expressing tumors.

## Introduction

Cell therapy with genetically engineered T cells expressing chimeric antigen receptors (CAR-T cells) specific for lineage antigens has shown therapeutic efficacy and received approval in a range of hematologic malignancies[1, 2]. Successful translation of CAR-T therapies to the treatment of patients with solid tumors has encountered several challenges, lack of tumor-specific cell surface antigens being one of them[3].

Compared with CARs, TCRs can address antigens from any tumor cell compartment including intracellularly expressed tumor-associated and tumor-specific antigens (TAAs, TSAs)[4–6]. Recent successful developments with TIL-therapies support this concept[7–9]. However, TCR recognition depends on peptide presentation by HLA molecules, which dictates that a therapeutic TCR can only be used in HLA-matched patients with antigen-positive tumors. As a result, most clinical TCR-T studies to date have focused on peptides from common TAAs presented by the highly prevalent HLA-A*02:01[10]. *Afami-cel* (*Afamitresgene autoleucel,* marketed as *Tecelra*) targeting MAGE-A4/HLA-A*02:01 in patients with sarcoma[11] and *Tebentafusp*, a TCR-derived bispecific receptor recognizing gp100/HLA-A*02:01 and CD3 approved for uveal melanoma are prominent examples[12]. TCR-T clinical trials targeting other shared epitopes of common TAAs have observed cases of severe on-target/off-tumor reactivity, as even low expression of TAAs in few normal tissues resulted in severe autoimmune side effects including fatal incidences with affinity-optimized TCRs against MAGE-A3[13–17].

In addition to toxicity risks, the TAA-directed TCR-T therapies described above are only available for a minority of patients. These barriers can be overcome by using natural TCRs to target TSAs, which include all types of somatic non-synonymous mutations in canonical proteins and aberrantly transcribed and translated gene products, collectively referred to as neoantigens[5]. While neoantigens were shown to be fundamental for the effects of immune checkpoint inhibition (ICI) and TIL therapy, only a small fraction was found immunogenic[18–23]. In addition, neoantigen expression is often heterogenous in tumors and metastases, and most neoepitopes are unique to individual tumors. Therapeutic exploitation requires personalization and identifying productive TCR-neoepitope-combinations is time- and labor-intensive[4, 24]. Nonetheless, personalized TCR-T cell therapy approaches targeting private neoepitopes in patients with refractory solid cancers are in clinical development [25, 26]. Neoepitopes derived from recurrent mutations in oncogenes are considered optimal targets because they drive tumorigenesis and progression, and exhibit clonal and stable expression across lesions[27]. Even though naturally occurring T cells specific for recurrent neoantigens have only occasionally been reported in patients[4, 27–29], their principal therapeutic activity has been demonstrated in the clinic[29–31]. Also, developments with synthetic immune receptors based on TCR-mimic antibodies recognizing prevalent oncogene peptide-HLA-complexes show the substantial interest in targeting these neoantigens[32–34]. TILs of individual cancer patients harbor polyclonal populations of tumor-specific T-cell clonotypes targeting private and shared tumor antigens, clonal driver mutations included. While they probably represent patient-specific optimal combinations of immunodominant T-cell responses they are mostly diluted in larger pools of tumor-nonspecific bystander T cells[35]. At least in part the tumor-experienced clonotypes are exhausted or dysfunctional reducing their responsiveness to current TIL expansion protocols[36].

A direct antigen-agnostic identification of tumor-specific T-cell clonotypes from TILs, sequencing and cloning of the most promising TCRs for manufacture of autologous TCR-T cells provides a treatment option for many patients. Current developments employ sorting of candidate tumor-specific T cells based on selective cell surface markers or single-cell gene expression signatures[37–43]. However, from a manufacturing and regulatory perspective, it is not clear as to whether these methods truly select only tumor-specific TCRs and how the most efficient ones are chosen for therapy. Personalized neoantigen-specific TCR-T approaches have shown that it is feasible to manufacture cell products with two to three different TCRs per patient[25, 26]. Similarly, for an antigen-agnostic TCR selection approach, it would be straightforward to select TCRs from a variety (2-4) of immunodominant anti-tumor clonotypes to address antigen heterogeneity and immune escape mechanisms.

In this study, we introduce an antigen-agnostic method to identify tumor-specific T-cell clonotypes by comparative high throughput TCR-repertoire profiling of tumor- and adjacent normal tissue-infiltrating T-cell clonotypes from surgical specimens. In seven NSCLC patients we identified candidate tumor-specific TIL clonotypes, in six of them the selection was supported by single-cell gene expression profiling. Experimental validation in three patients revealed that tumor-specific clonotypes predicted by our method responded to autologous tumor cells. For one of these patients, we simulated the production of therapeutic TCR-T cells by selecting four tumor-specific TIL clonotypes, decoded their αβTCR sequences, synthesized and expressed them in healthy donor T cells. Screening with the TCR-T cells for recognition of expressed non-synonymous neoepitopes and overexpressed TAA candidates revealed that three of the four TCRs specifically recognized mutant KRAS Q61H-peptide ILDTAG**<u>H</u>**EEY presented by HLA-A*01:01. The tumor-specificity and therapeutic potential of the selected TCRs are reinforced by functional characterization of the TCR-T cells, the gene expression signatures of the original TIL clonotypes, the fact, that the clonotypes were found infiltrating a tumor relapse acquired more than 30 months after surgery of the primary tumor, and the discovery of highly homologous to identical TCRs in six of 29 archival (FFPE) tumor samples with confirmed KRAS Q61H-mutation. The results highlight our method’s capacity to directly select tumor-specific TCRs for therapy and suggest the mutant KRAS-specific TCRs as candidates for an off-the-shelf TCR-T therapy in HLA-A*01:01-positive patients with RAS Q61H-positive tumors.

## Results

### Clinical data of NSCLC patients and acquisition of clinical materials

For seven patients, fresh tumor and adjacent normal lung specimens were selected by pathologist from excess material and transported to the laboratory along with a peripheral blood sample for immediate processing. The disposition of the study patients with respect to the experimental strategy is shown in **Figure 1A**. All underwent lobectomy and lymph node dissection with curative intent. In three patients, functional analyses were performed: patient 1 (m/57) was diagnosed with lung squamous carcinoma of the left lower lobe in May 2016, patient 2 (f/73) with adenocarcinoma of the right superior/middle lobe in September 2020, and patient 3 (f/54) with lung adenocarcinoma of the left superior lobe in June 2018. For patient 3, in addition to clinical samples from surgery of the primary tumor, follow-up samples including a tumor recurrence, blood and plasma samples were analyzed. Hence, more details on her clinical course and the derivation of the clinical specimens analyzed are provided: In July 2019 a local recurrence was diagnosed via PET-CT and the patient received concomitant chemoradiotherapy followed by durvalumab maintenance therapy for one year. In January 2021 the local recurrence localized in the aortopulmonary window increased in size. An extended pneumonectomy was performed. From the recurrent tumor, formaldehyde-fixed paraffin-embedded (FFPE) tissue samples were preserved by pathologist. Further blood samples were collected and processed in September and December 2021. The patient’s clinical course is summarized in **Table S1** and sampling-time points are given in **Fig. S1**. Immunohistochemistry (IHC) of consecutive FFPE slices revealed that patient 3’s primary and recurrent tumors were positive for HLA-A expression and showed a PD-L1 proportion score >50% (**Fig. S2C,D**). CD3-, CD4-, and CD8-positive TIL subpopulations in the primary and in the recurrent tumor were prevalent in peritumoral areas rather than in the tumor core (**Fig. S2A,B**). The patient has been in sustained clinical remission since 2021.

**Figure 1:**
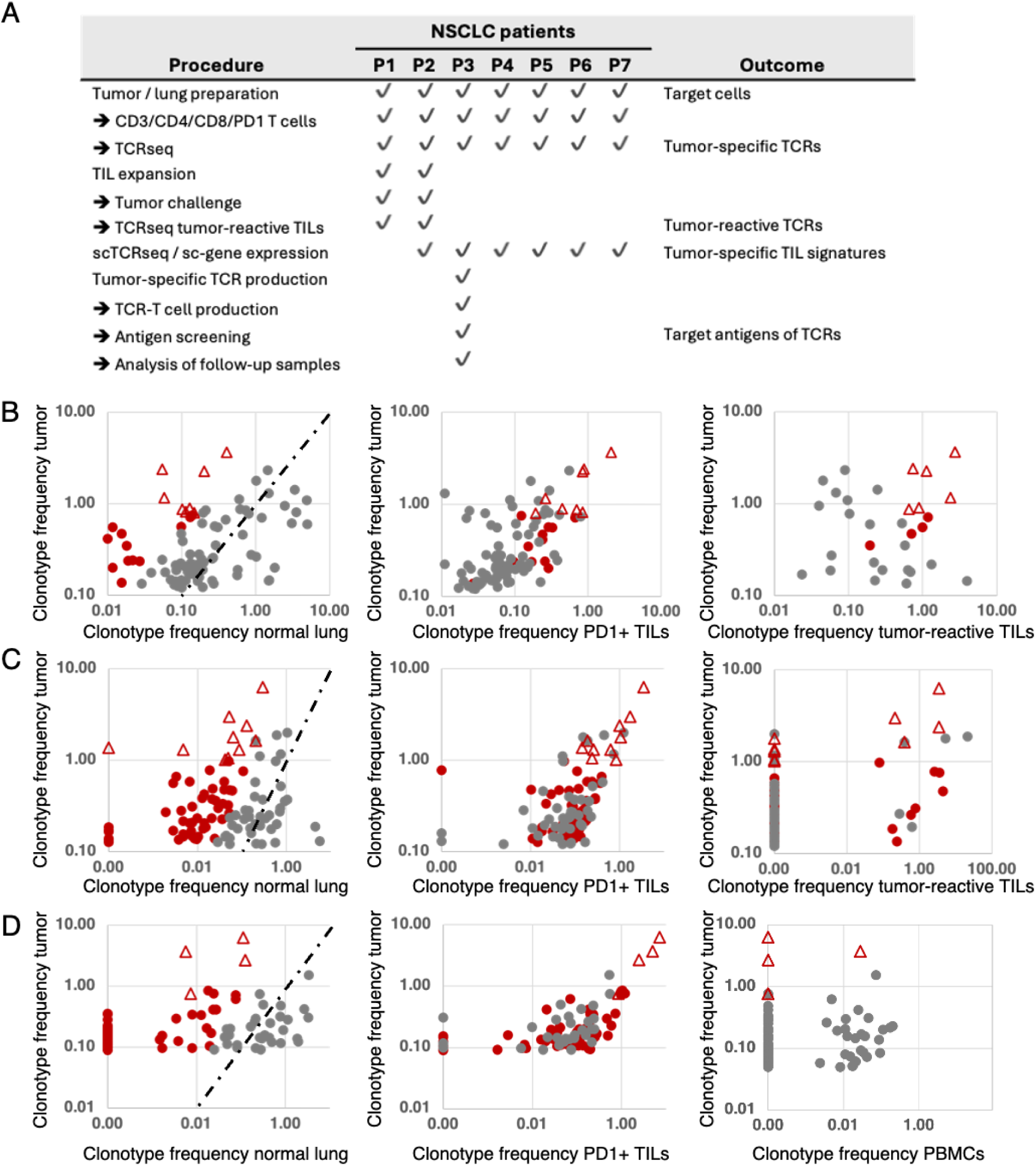
Disposition of the study patients (A) and identification and selection of tumor-reactive T-cell clonotypes in patients 1 (B), 2 (C) and 3 (D). Frequencies of T-cell clonotypes (percentages) were determined by TCRseq from TILs, adjacent normal lung, the PD1-positive fraction of TILs and (patient 1, patient 2) after TIL-expansion and stimulation with autologous tumor cells. The top 100 TIL clonotypes of each patient were analyzed in relation to their frequency in normal lung (left), in the PD-1-positive TIL fraction (middle), and in the tumor-reactive fraction after stimulation with autologous tumor cells (B,C, right). Tumor-specific clonotypes were predicted according to high TIL-frequency and a frequency ratio >5 resulting from comparing the frequencies of TIL- and normal lung-infiltrating clonotypes (tumor-to-nontumor ratio, B-D, left). Clonotypes with ratios >5 are depicted in red. The dashed lines indicate ratio=1 values. Red triangles represent clonotypes selected as best candidates for tumor-specific T cells and possible sources of therapeutic TCRs. For patient 1, eight clonotypes were initially selected (B, left). The same clonotypes were highly abundant among PD1-positive TILs (B, middle). After expansion in vitro, stimulation with autologous tumor cells and sorting by IFN-γ-capture assay, six of the eight selected clonotypes showed tumor reactivity (B, right). For patient 2, ten clonotypes were initially selected (C, left) and shown expanded among PD1-positive TILs, too (C, middle). After in vitro expansion, stimulation with autologous tumor cells and sorting for CD137-positive cells, four of the ten selected clonotypes were found to be tumor-reactive (C, right). For patient 3, no in vitro expansion of TILs was performed. Instead, the TCRs of the top four clonotypes according to TIL-frequency and high tumor-to-nontumor ratio (D, left) and high frequency among PD—positive TILs (D, middle) were selected, subjected to scTCRseq, synthesis and cloning. The four recombinant TCRs were used to produce TCR-T cells to show their tumor-reactivity and apply them to the screening of target antigens the TCRs recognize.

### Preparation of surgical material and identification of tumor-specific TILs

Primary tumor and adjacent lung tissues, as separated by pathologist, were physically and enzymatically dissociated and from the resulting cell suspensions as well as blood-derived PBMCs, CD3-, CD4-, CD8-positive lymphocyte fractions were sorted (**Figure 1A**). PD-1-positive lymphocytes were sorted from TILs. The remains of the tissue cell suspensions were cryopreserved. Genomic (g)DNA isolated from all T-cell fractions was used as template for TCR-VDJ-amplification and sequencing (TCRseq) to profile the αβTCR-repertoires of all TIL- and lung T-cell fractions as described in [44, 45] (**Figure 1A**). As exemplified for patients 1-3, candidate tumor-specific T-cell clonotypes were determined by comparing the frequencies of tumor-infiltrating with lung-infiltrating clonotypes. Based on high tumor prevalence and a tumor-specific distribution (frequency ratio tumor-to-nontumor >5), CD8-positive clonotypes were predicted as candidate tumor-specific T cells (**Fig. 1B,C,D left graphs**). High frequencies of candidate clonotypes in PD-1-positive TIL fractions supported the selection (**Fig. 1B,C,D middle**). In patients 1 and 2, TILs were subjected to in vitro expansion using an in-house protocol (patient 1, **Fig. S3**) or a small-scale rapid expansion protocol adapted from a clinical TIL manufacturing protocol (patient 2, **Fig. S4**)[46]. After two to three weeks of culture, expanded TILs were challenged with autologous tumor cells and sorted via IFN-γ Secretion Assay – Cell Enrichment and Detection Kit (Miltenyi, patient 1, **Fig. S3**) or CD137-expression per FACS (patient 2, **Fig. S4**). Tumor-activated IFN-γ- and CD137-positive cells were subjected to TCRseq, their frequencies were determined and compared to the top 100 TIL clonotypes at the starting time point (**Fig. 1B,C right**). In patient 1, six of eight candidate tumor-specific clonotypes showed tumor-reactivity (**Table S2, Fig. 1B right**), in patient 2, four of ten predicted tumor-specific clonotypes responded to tumor challenge (**Table S3, Fig. 1C right**). In both patients, the tumor-reactive clonotypes represented the top six (patient 1, **Table S2**) and top four (patient 2, **Table S3**) clonotype candidates determined before. Having shown that our method can predict tumor-specific clonotypes based on comparative TCRseq between TILs and normal tissue-resident T cells, we set out to analyze the TCRs of the top four predicted tumor-specific clonotypes of patient 3 using a TCR-T cell approach. As for patients 1 and 2, the TCR selection for patient 3 was based on TIL-prevalence, high tumor-to-nontumor frequency ratio and high frequency among PD-1-positive TILs (**Table S4, Fig. 1D**). The selected TCRs were designated TCR-V1,-V2,-V3, and -V4 (**Fig. 2A**) and alignment of the TCRs’ CDR3 sequences revealed striking sequence homologies between TCR-V1, -V2, and -V3 suggesting a shared antigen specificity (**Fig. 2A**). The predominance in the tumor was not reflected in peripheral blood, as three of the four selected clonotypes were absent from blood lymphocytes, one was detected only at low frequency (TCR-V2, 0,02%, **Fig. 1D, right**). Single-cell RNA sequencing (scRNA-Seq) of TILs decoded the paired αβTCR chains of selected clonotypes (**Table S5**), and sc-gene expression profiling revealed functional properties and differentiation trajectories of the cells (see below). The TCRs were synthesized as bicistronic chimerized expression constructs (cTCRs) with the human constant domains of the chains replaced by murine homologs (**Fig. 2A**) and cloned into vector pMX-puro for retroviral transduction of human T cells from healthy donors.

**Figure 2:**
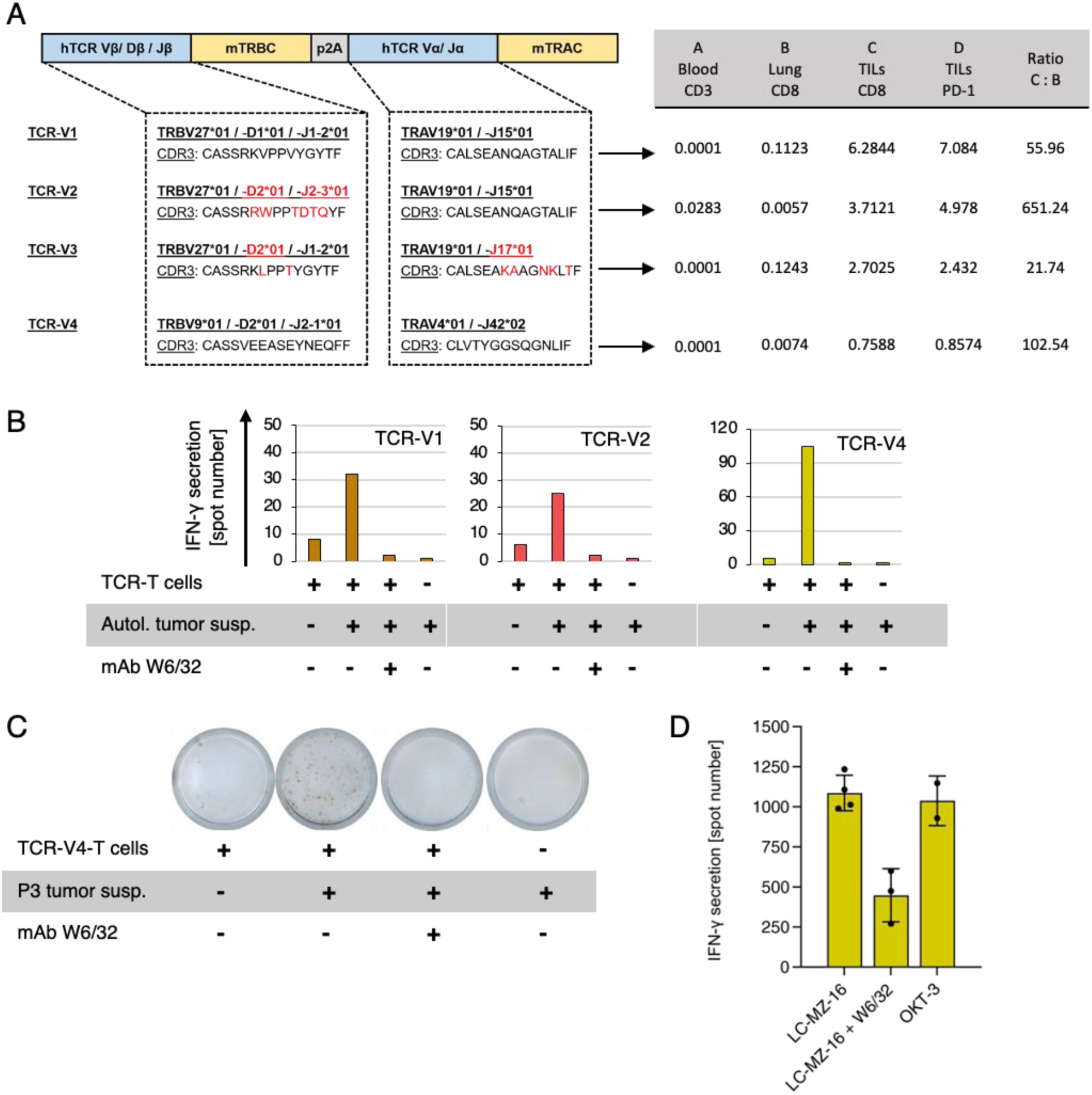
: (A) Schematic representation of the patient 3-TCR constructs synthesized and cloned for functional analyses. Human constant domains were replaced by murine homologous sequences. V-(D)-J gene segments of α- and β-chains and CDR3 sequences are shown. Red labels highlight differences between TCR-V1, -V2, and V3. Key data leading to the selection of the four T cell clonotypes for functional characterization are detailed in the adjacent table. Production, characterization and tumor-response analysis of recombinant P3-TCR-T cells. (B) Recognition of patient 3 tumor cells by TCR-T cells as determined by IFN-γ ELISpot assay. Tumor cell suspension after thawing was sufficient for only one experiment. (C) Original ELISpot well-scans showing the tumor-response of TCR-V4-T cells. (D) Recognition of HLA-A2-matched allogeneic NSCLC cell line LC-MZ-16 by TCR-V4-T cells (summary of four independent experiments). OKT-3 mAb was used for unspecific activation. Pan-HLA class I mAb W6/32 blocked tumor-recognition in all experiments.

### Production and functional characterization of tumor-specific TCR-T cells

Tumor-specific TCR-T cells were produced by retroviral transduction of donor-derived T cells with synthesized codon-optimized sequences encoding TCR-V1-V4 (**Fig. 2A**) as described[47]. Before retroviral transduction, the recipient T cells were depleted from endogenous TCRs by CRISPR/CAS9-mediated knock-out (KO) of human (h)TRBC and (h)TRAC domains to prevent mispairing of endogenous and recombinant TCR chains with unpredictable adverse specificities or allo-reactivity against allogeneic antigen-presenting cells used for subsequent antigen-screening. Successful hTCR-KO and expression of recombinant cTCRs was confirmed by flow cytometry (**Fig. S5**). IFN-γ-ELISpot-assays with the cTCR-T cells showed recognition of patient 3’s tumor cells (**Fig. 2B,C**). When all TCR-T cells were tested against HLA-matched tumor cell lines (not shown), only TCR-V4 T cells responded by recognizing the HLA-A*02:01-matched lung cancer cell line MZ-LC-16 (**Fig. 2D**). Because HLA-A*02:01 is the only allele matched between both tumors, this finding indicated HLA-A02-restriction of TCR-V4 and suggested expression of a target antigen shared between patient 3 tumor cells and MZ-LC-16. MHC class I restriction of all four TCRs tested was demonstrated by blockade with the pan-HLA class I antibody W6/32 (**Fig. 2B,C,D**).

### Target antigen screening using tumor-specific TCR-T cells

Neoantigens have been associated with favorable clinical responses to immunotherapy in NSCLC. Therefore, comparative whole exome- and transcriptome sequencing of tumor and adjacent lung tissue samples were carried out to identify tumor-specific non-synonymous variants as neoantigen candidates (**Fig. S6**). Seventy-three expressed non-synonymous single nucleotide variants (SNV) and one frameshift-mutation were identified (**Fig. S6, Table S6**). Structural variant analysis revealed no translocations distinctive of subtypes of NSCLC (not shown). Binding predictions of mutated candidate peptides to the patient’s HLA I alleles (HLA-A*01:01/*02:01, HLA-B*08:01/*40:02, HLA-C*03:04/*07:01) using IEDB (http://tools.iedb.org/mhci/) and NetMHC4.0 (https://services.healthtech.dtu.dk/service.php?NetMHC-4.0) public databases found 581 9- or 10-mer peptides with IC50 <500nM and/or percentile rank <6 (**Table S7**). HLA allele-assorted peptides were affinity score-ranked, and the 94 top-scoring peptides (and two quality control peptides) were synthesized and tested for recognition (**Table S7**). K562 cells transduced with any of the patient’s HLA I alleles were pulsed with candidate peptides and tested for recognition by TCR-T cells using ELISpot assays. TCR-T cells expressing any of TCR-V1, -V2 and -V3 responded to KRAS Q61H peptide 55-64 ILDTAG**H**EEY, regardless of whether CD4- or CD8-positive lymphocytes expressed the TCRs. (**Fig. 3A,B**). TCR-V4-T cells failed to recognize any of the mutated peptides tested. Because TCR-V4-T cells were shown before to respond to stimulation with NSCLC line MZ-LC-16 (**Fig. 2E**), we suspected a target epitope shared between patient 3’s tumor and the cell line. Comparative analysis of non-synonymous variants as determined by WES found no shared mutated neoantigen in both tumors (**Fig. S7A**). Differential gene expression analysis of tumor versus normal lung tissues revealed overexpressed transcripts in both tumor entities (**Fig. S7B,C,D**). Shared overexpression was found only for cancer-germline antigens CT83, MAGEA12 and XAGEA1. TCR-V4-T cells were tested against 293T cells co-transfected with antigen- and HLA-A*02:01-coding cDNAs by ELISpot. However, none of the three candidates was recognized (**Fig. S7E**) and the cognate antigen of tumor-specific TCR-V4 was not found.

**Figure 3:**
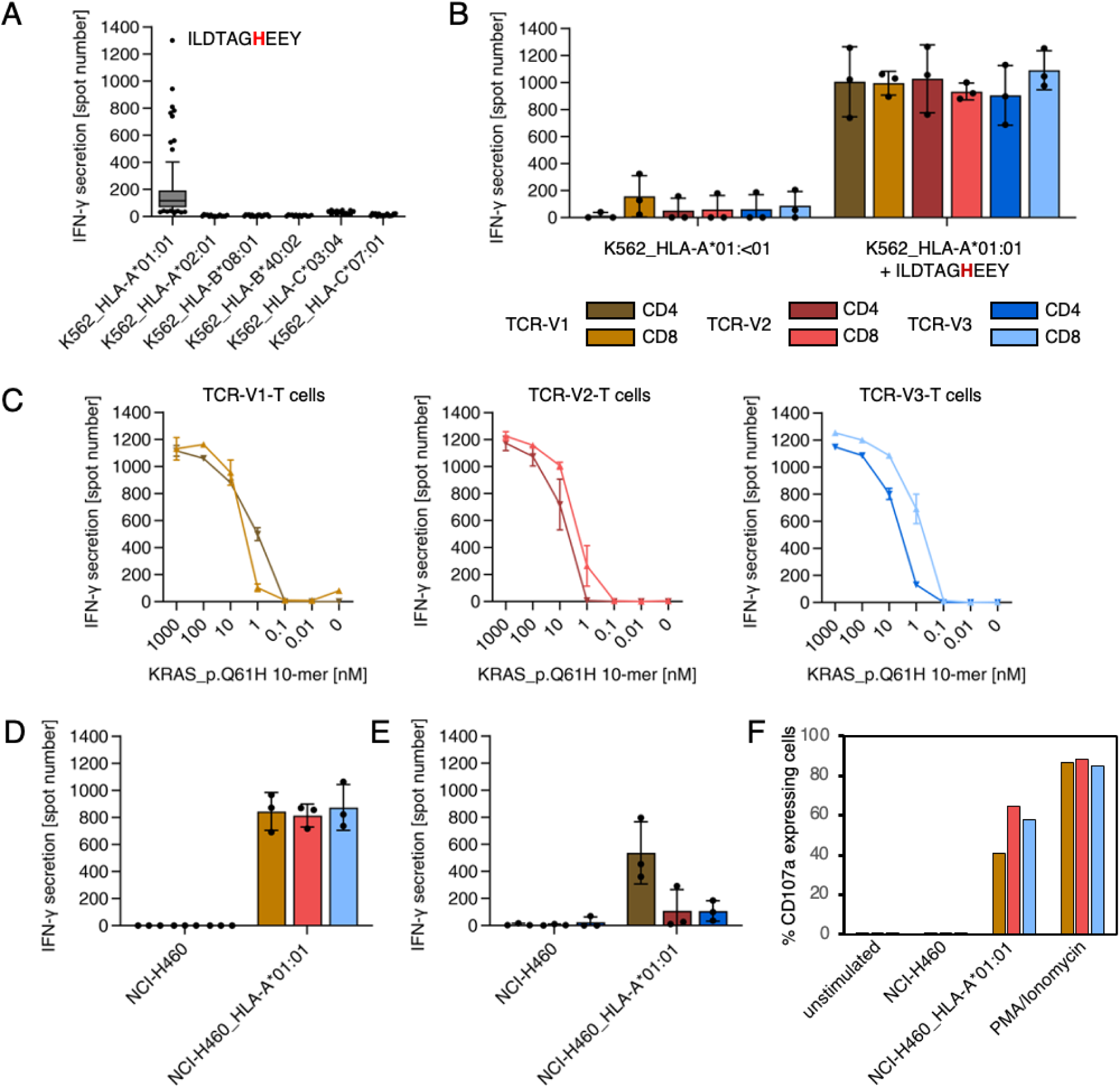
TCR-T cells transduced with P3-TCRs V1, V2, V3 recognize the naturally processed and presented KRAS Q61H-peptide 55-64 (ILDTAGHEEY). (A) Identification of the KRAS Q61H-peptide as target antigen of TCR-V2-T cells. ELISpot analysis testing TCR-T cells against monoallelic P3-HLA transduced K562 cells pulsed with 96 candidate neoantigen peptides identified by WES- and RNA-Seq. Only K562/HLA-A*01:01 cells were recognized when pulsed with several synthetic peptides in a cross-reactive manner, though less strong as the KRAS Q61H-peptide. (B) K562/HLA-A*01:01 cells were recognized by TCR-V1, -V2, and –V3-transduced CD4- and CD8-positive TCR-T cells when pulsed with mutant KRAS-peptide 55-64 (color code of the legend used for all figures). (C) K562/HLA-A*01:01 cells pulsed with titrated doses ILDTAGHEEY were recognized by TCR-T cells with high functional avidity (EC50<10nM). (D,E) Recognition of KRAS Q61H-mutated, HLA-A*01:01-transduced cell line NCI-H460/HLA-A*01:01 in comparison to wildtype NCI-H460 by CD8-positive (D) and CD4-positive (E) TCR-T cells. (F) Flow cytometry showing degranulation (CD107a) as surrogate for lytic activity of TCR-V1, -V2, and -V3-expressing CD8-positive TCR-T cells upon co-culture with NCI-H460/HLA-A*01:01. Corresponding results for CD4-positive TCR-T cells and the experiment gating strategy are shown in Fig. S7. Lytic activity and cytokine release of all TCR-T cell cultures showed overlapping results. All ELISpot experiments were done in duplicates or triplicates. Results shown in B,D,E are derived from three independent experiments.

### Characterization of the three distinct KRAS Q61H-reactive TCRs

For a more comprehensive analysis of the three mutant (m)KRAS-specific TCRs, CD4- and CD8-positive T cells transduced with TCR-V1, -V2, or -V3 were tested against K562/HLA-A*01:01 cells pulsed with titrated doses of the mKRAS peptide 55-64. All TCR-T cells showed recognition at EC_50_ values below 10nM regardless of whether the three TCRs were expressed in CD4- or CD8-positive TCR-T cells (**Fig. 3C**). To verify that the mKRAS peptide is processed and presented, NCI-H460 cells were tested for recognition. NCI-H460 cells are natural carriers of the KRAS Q61H-encoding mutation (KRAS c.183A>T) but are negative for HLA-A*01:01 and were thus transduced with this allele. Wildtype NCI-H460 and NCI-H460/HLA-A*01:01 cells were tested for TCR-T-cell recognition by ELISpot (**Fig. 3D,E**) and degranulation as a surrogate assay for cytolytic activity (**Fig. S8**). While NCI-H460 cells induced no response, as expected, NCI-H460/HLA-A*01:01 cells were strongly recognized by CD8-positive TCR-T cells transduced with any of the three TCRs (**Fig. 3D,F; Fig. S8B,C**). In contrast, CD4-positive TCR-T cells showed significant reactivity only when transduced with TCR-V1 (**Fig. 3E, Fig. S8C**). Weaker responses of TCR-V2 and -V3-transduced CD4-positive T cells against NCI-H460/HLA-A*01:01 suggest dependency on CD8-costimulation of TCR-T cells transduced with these TCRs. H-, K-, and NRAS protein-family members share identical Aa sequences from position 1 to 86 (**Fig. S9A**) implying that the peptide comprising Aa 55-64 can be processed and presented from any of these proteins. Prevalent alterations at the RAS mutation hotspot Aa position 61 include Q61H, Q61R, Q61K and Q61L, which occur with different frequencies in different tumor entities (**Fig. S9B**). Taking advantage of three independently evolved homologous but not identical KRAS Q61H-specific TCRs (**Fig. 2A**), we tested whether any of the TCRs was capable of cross-reacting against one of the alternative mutations. ELISpot assays with 293T cells transfected with HLA-A*01:01 and cDNAs encoding all four possible mKRAS variants as well as wildtype KRAS showed that all three TCRs are specific for the Q61H mutation (**Fig. 4A**). As a definite proof that KRAS Q61H is the actual target of the TCRs, CRISPR/CAS9 technology was used to change the Q61H-encoding mutation in NCI-H460/HLA-A*01:01 cells to encode KRAS Q61R (codon alteration KRAS c.181-183CAT>CGC; **Fig. S10**). Because NCI-H460 cells are homozygous for the mutation, both alleles had to be edited to achieve an effect on TCR-T cell recognition. Treated tumor cells were cloned by limiting dilution and, after expansion, multiple clones were tested for recognition by the TCR-T cells. Patterns of recognition observed included unaltered, reduced and lost recognition. Sequencing of target genomic regions of one representative tumor clone for each pattern revealed that TCR-T cell recognition correlated with the extent of target codon editing (**Fig. 4B**): failed editing resulted in unaltered recognition (clone 9), conservation of only one of the two Q61H-encoding alleles reduced recognition (clone 11), and the successful biallelic codon editing encoding KRAS Q61R resulted in loss of recognition (**Fig. 4B**). The results demonstrate that all three TCRs only recognize NCI-H460/HLA-A*01:01 cells expressing the KRAS Q61H neoepitope but none of the alternative hotspot-neoepitopes, suggesting a strict target-specificity. Concerning cross reactivity, in addition to the lack of responses to the related peptides, the TCR-T cells did not respond to the various APCs used in different assays, including transfectants expressing the patient’s HLA alleles, involving K562 cells, 293T cells, and an HLA-matched lymphoblastoid cell line (not shown). Furthermore, a search with the cognate target peptide of the CrossDome database[48] for processed and presented peptides from normal tissues did not yield any peptide hits with cross-reactive potential (not shown).

**Figure 4:**
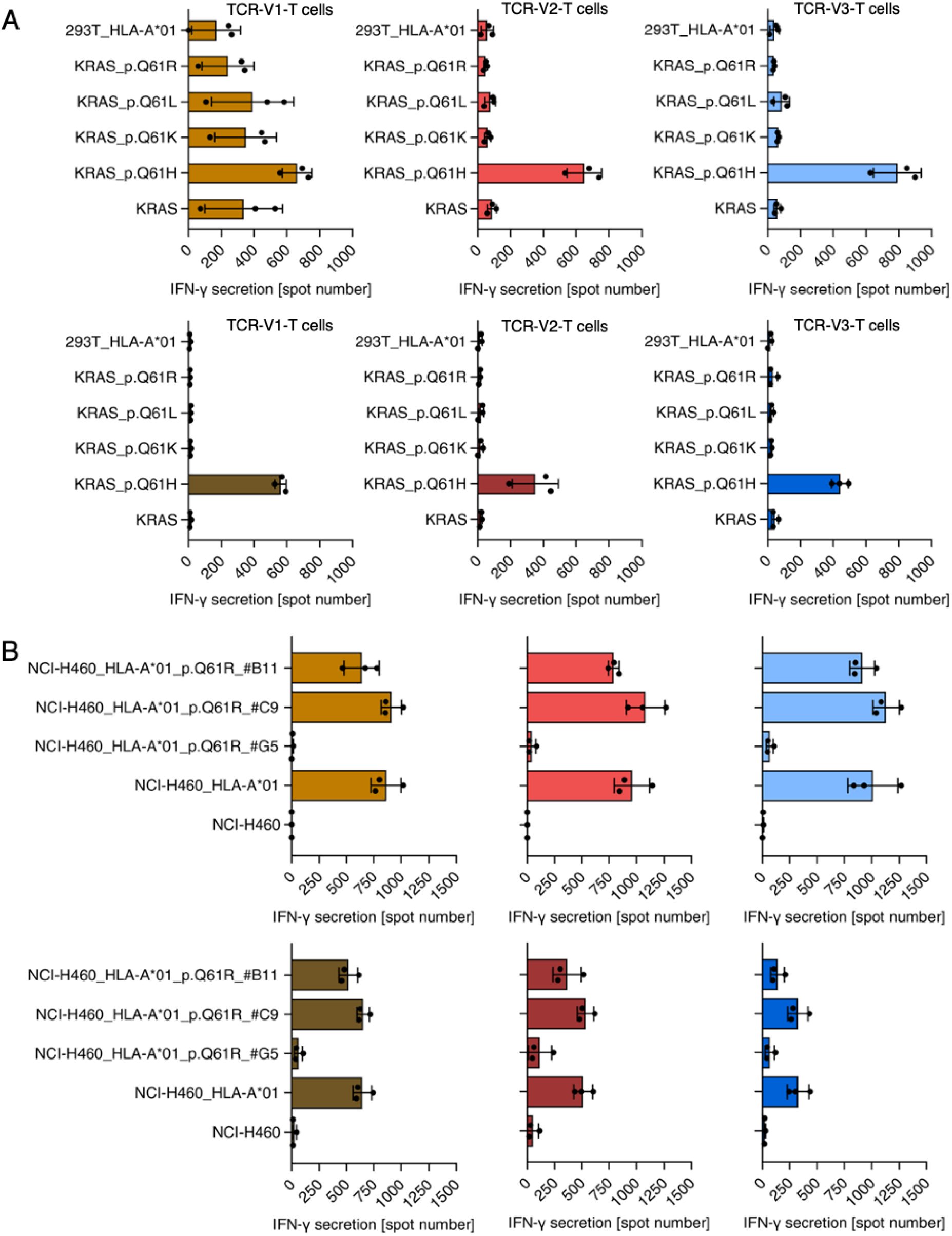
TCR-T cells transduced with TCRs V1, V2, V3 are KRAS Q61H-specific. (A) Reactivity of TCR-V1/V2/V3 transduced CD8-positive (top row) and CD4-positive (bottom row) TCR-T cells against 293T cells transiently transfected with depicted KRAS-encoding variants and HLA-A*01:01. TCR-V1-transduced CD8-positive T cells produced comparable background activity against 293T cells expressing KRAS-wt, KRAS Q61L, Q61K, and Q61R. Recognition of KRAS Q61H was stronger. All other TCR-transduced CD8- and CD4-positive TCR-T cells showed exclusive recognition of KRAS Q61H-/HLA-A*01:01-expressing 293T cells. (B) Reactivity of TCR-V1/V2/V3 transduced CD8-(top) and CD4-positive (bottom) TCR-T cells against NCI-H460, NCI-H460/HLA-A*01:01, and NCI-H460/HLA-A*01:01 cells treated by CRISPR/CAS9 for replacement of the Q61H-encoding mutation by Q61R-encoding sequences. Treated cells were cloned by limiting dilution, expanded and after target region sequencing tested for recognition by CD8- and CD4-positive TCR-T cells. Three examples with different outcomes are shown. NCI-H460/HLA-A*01:01 clone #G5 carries a biallelic substitution encoding KRAS Q61R and recognition of clone #G5 was lost. Clone #B11 harbors a frameshift mutation in one of two KRAS Q61H alleles, explaining the reduced recognition of the cells. In #C9, the H-to-R substitution failed explaining the retained recognition of the cells. All ELISpot experiments were done in duplicates or triplicates. Shown are results of three independent experiments.

### Course of the KRAS Q61H response in the patient over time and presence of matching TCRs in independent KRAS Q61H-positive tumors

Consistent with KRAS Q61H being a cancer driver in NSCLC, the hotspot variant was clonal and detected in genomic DNA from a tumor relapse obtained 32 months after surgery of the primary tumor (**Table S1, Fig. S1**). TCRseq using template DNA from the relapse-FFPE sample detected multiple clonotypes predicted tumor-specific from the primary tumor, including all four confirmed tumor-reactive TCRs (TCR-V1/V2/V3/V4) at highest frequencies (**Fig. 5A**). Moreover, the TCR-V1-coding sequence was detected by TCR-repertoire sequencing from plasma cfDNA from a blood sample collected in September 2021 (**Fig. S1**; **Fig. 5B**) suggesting cellular turnover of this clonotype at this time point *in vivo*. By contrast, the clonotype was undetectable in PBMCs from blood collected at the same timepoint and three months later.

**Figure 5:**
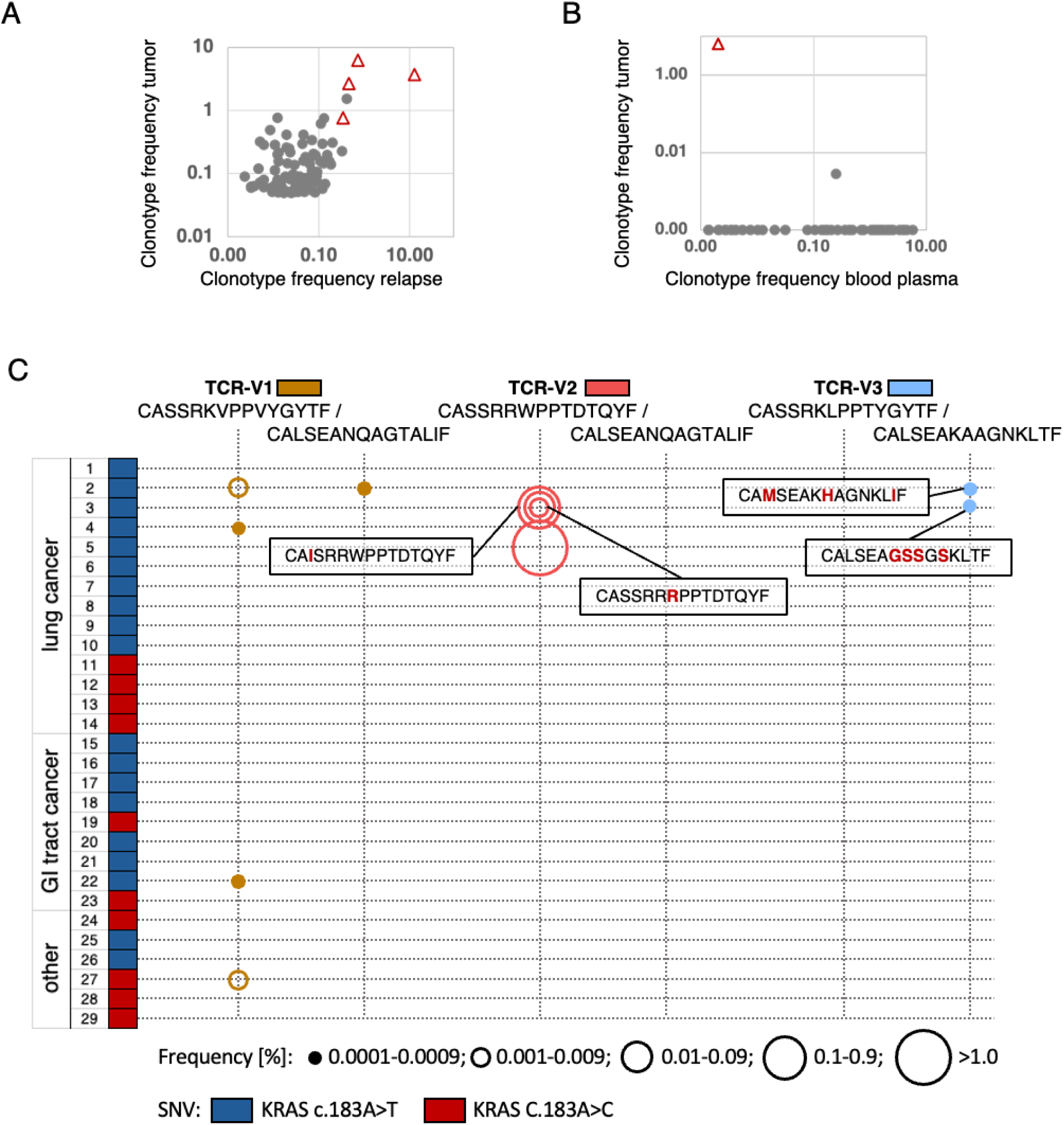
TCR-V1–V4-positive clonotypes infiltrating the P3 relapse tumor, detected in plasma and homologous TCRs in archival tumor samples of independent patients. (A) Frequencies of the four selected clonotypes (triangles) amid the top 100 clonotypes detected in primary tumor and relapse. (B) Frequency of the TCR-V1-clonotype (triangle) among TCR sequences amplified from plasma-derived cfDNA of P3. The top 100 primary tumor clonotypes were compared with clonotypes detectable in a plasma sample from September 2021. In corresponding blood from the same and a later time point, none of the clonotypes was detected. (C) Discovery of TRBV/TRAV-CDR3 sequences matching to the mutated KRAS-specific P3-TCRs in samples from patients with KRAS Q61H-positive tumors (29 FFPE samples tested, mutations encoded by c.183A>T or c.183A>C are symbolized by color). Sequence-identical or highly related TCRs were found in 6/29 patients. Only non-identical CDR3-sequences are posted, Aa differences highlighted in red. TCR-frequency ranges are represented by circle sizes.

To further investigate the immunogenicity of the KRAS Q61H mutation, we performed TCRseq on gDNA from FFPE-samples of various KRAS Q61H-positive tumors from 29 patients including 14 lung cancers, nine gastro-intestinal (including CRCs, pancreatic, and bile duct cancers), and six not otherwise specified tumors (others, **Fig. 5C, Table S8**). The KRAS-mutation was encoded by SNVs c.183A>T in 19 and c.183A>C in ten cases. Repertoire-analysis of TRBV- and TRAV-sequences revealed CDR3 sequences highly related or even identical to patient 3 TCR-V1, -V2, and -V3 in six out of the 29 patient samples analyzed (in 4/14 lung cancers). Most frequently detected was the exact TRBV-sequence of TCR-V1 in four samples (2 lung, 1 rectum and 1 other cancer **Fig. 5C, Table S8**). In lung cancer sample-2, in addition to the matching beta the alpha-chain of TCR-V1 was found. This sample contained also an alpha chain related to TCR-V3 (**Fig. 5C, Table S8**). In FFPE-samples 3 and 5, multiple TCR-V2-matching TRBV-sequences were discovered. However, being aware that sequencing of DNA/RNA from paraffin-material is riddled with artifacts[48], we considered only perfect matches and sequences represented in elevated frequencies (>0.001%, coverage >4reads) as true hits in these cases (**Fig. 5C, Table S8)**. Of note, in sample-5 a TCR-V2 beta-chain perfect match represented 2.9% of all detected clonotypes. Taken together, these results suggest a convergent selection of cognate immune receptors in different patients with KRAS Q61H-positive cancers, suggesting a high epitope immunogenicity.

### scRNA-Seq of TILs reveals differentiation trajectories of tumor-specific T-cell clonotypes consistent with cytotoxicity, chronic stimulation and exhaustion

Single-cell gene expression analysis can inform about activation and differentiation states of TIL clonotypes. We analyzed a pool of about 13,000 single T cells from six NSCLC patients including patients 2 and 3 (**Fig. 1A**). To select tumor-specific clonotype candidates, a rigorous threshold (tumor-to-nontumor ratio >10, absolute frequency of CD8-positive TIL >0.2%) was applied. Of all TILs analyzed, 160 clonotypes (830 single cells, 6,4%) were confirmed or predicted to be tumor-specific by our method. Unsupervised clustering of all cells separated five clusters of CD8-positive from five CD4-positive T-cell clusters (**Fig. 6A, B**). Following subclustering of only CD8-positive T cells (**Fig. 6C**, ≈6600 cells) we identified the predicted tumor-specific clonotypes, including the confirmed tumor- and KRAS Q61H-specific clonotypes from patients 2 and 3, in two of the resulting five clusters (clusters 0 and 2; **Fig. 6C, D**). Specifically, CD8-positive T cells in cluster 2 were enriched for genes associated with activation, cytotoxicity, and tissue homing (granzymes, IFNG, CXCL13, CXCR6) but also terminal differentiation and exhaustion (LAYN, TOX, PDCD1, HAVCR2, ENTPD1, etc.; **Fig. 6D,F**). As a control, predicted expanded bystander T-cell clonotypes (tumor-to-nontumor ratio <1, frequency >0.1%) were localized by barcodes (**Fig. 6E**) and were found mainly in clusters 1 and 4 – were scarce in cluster 0 and largely absent from cluster 2. Detailed analysis of single cells of tumor-specific clonotypes revealed differentiation trajectories ranging from effector-memory/resident memory to terminally differentiated/exhausted T cells, suggesting that the T cells have been activated by tumor cells and eventually became exhausted due to chronic antigen stimulation. Hence, for the top clonotypes selected based on large frequency, a high tumor-to-nontumor frequency ratio and PD-1-expression, their gene signatures indicating exhausted/dysfunctional T cells supported the selection.

**Figure 6:**
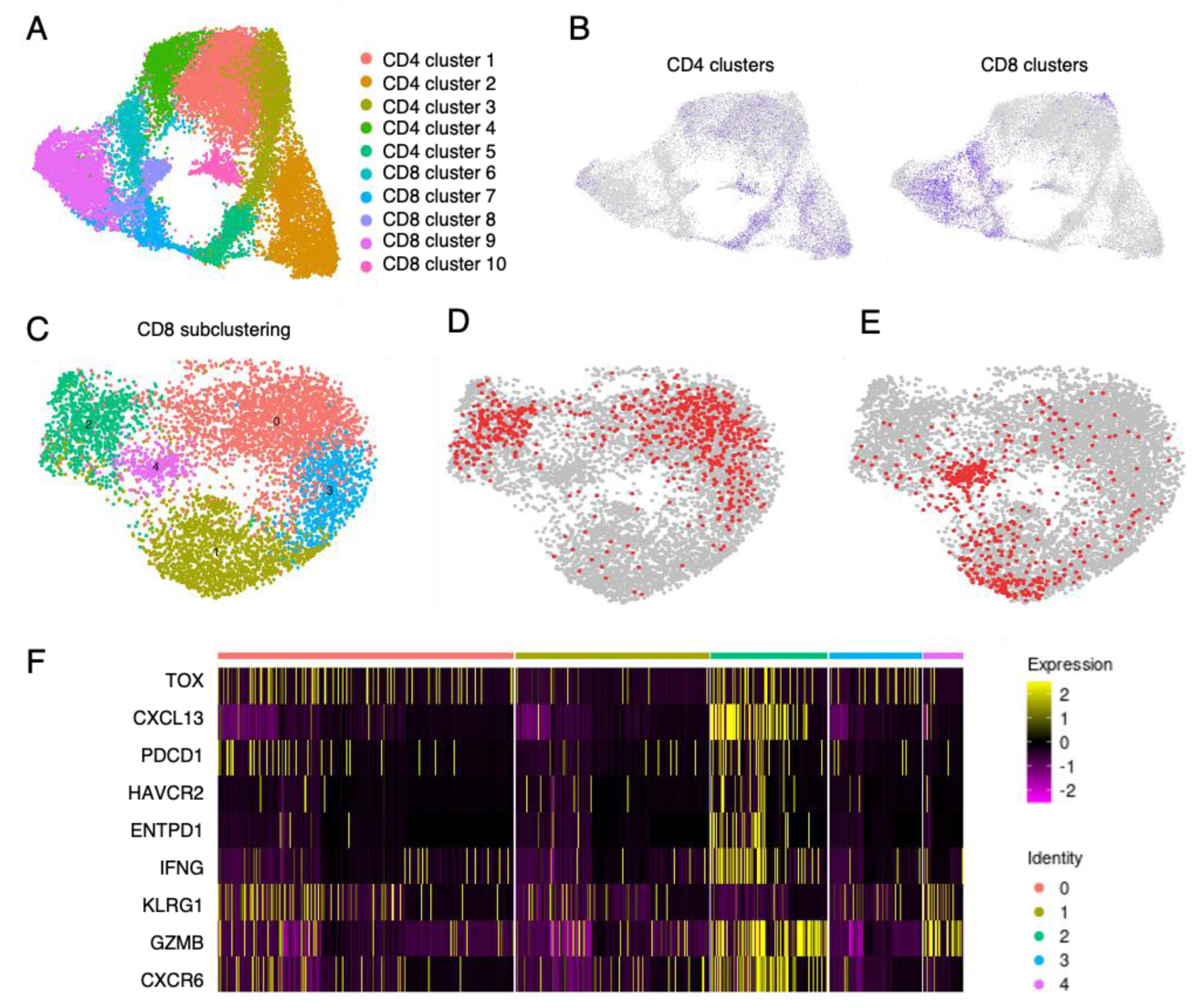
Single-cell gene expression analysis of TILs from six NSCLC patients coupled with barcode-mediated detection of predicted and confirmed tumor specific clonotypes. (A) Uniform Manifold Approximation and Projection (UMAP) of an unsupervised clustering of TILs from patient samples displaying five clusters each of CD4- and CD8-positive clonotypes. (B) Separation of CD4- and CD8-positive TIL clusters. (C) Subclustering of the CD8-clusters from B projects five new clusters. (D) Barcode localization of predicted (frequency >0.2%, tumor-to-nontumor frequency ratio >10) and confirmed tumor-specific clonotypes (in red). (E) Barcode localization of bystander clonotypes (frequency >0.1%, tumor-to-nontumor frequency ratio <1, in red). (F) Heatmap showing expression of genes associated with tissue residency, effector function and terminal differentiation/exhaustion. It is obvious that tumor-specific TILs (prevalent in clusters 0 and 2) and bystander T cells (concentrated in clusters 1 and 4) express marker genes of cytotoxicity and tissue residency. However, expression of exhaustion markers is high in clusters 0 and 2 and absent from clusters 1 and 4, efficiently separating bystander T cells from tumor-specific TILs.

## Discussion

Adoptive cell therapy with engineered tumor-reactive TCR-T cells expands the cellular therapy options for solid cancers. Compared with CARs, TCRs address a significantly larger antigen repertoire and TCR-T cells can recognize target epitopes with superior sensitivity[49, 50]. Higher functional avidity of TCR-T cells endows them with stronger tumor cell killing efficacy. At the same time, lower target-binding affinity enables serial scanning and killing of multiple tumor cells. In a therapeutic setting, this quality may delay exhaustion and increase persistence of the TCR-T cells[10]. Current TCR-T approaches in clinical development have in common that they are driven by an antigen-centered perspective. Either they target a very restricted number of antigens in combination with few common HLAs, mainly HLA-A*02:01[10] or, in a personalized approach, they focus on TCRs against private neoantigens [25, 26]. Both strategies are limited to small numbers of eligible patients.

In this study, we present an antigen-agnostic method to identify tumor-specific T-cell clonotypes based on (1) numerical dominance among TILs, (2) high tumor-to-non-tumor frequency ratios, (3) PD-1 expression, and (4) verification of clonotype selection by single-cell gene expression data showing that candidate clonotypes express gene signatures indicative of chronic activation, terminal differentiation and/or exhaustion. Compared to competing studies that have suggested the identification of tumor-specific T cells from TILs or peripheral blood using only single cell expression profiling [38–43, 51], our scoring matrix facilitates the direct selection of therapeutic TCRs from immunodominant clonotypes. A selection based on a combination of four qualifiers may be more likely to convince regulatory authorities to approve the testing of tumor-specific TCR candidates in clinical trials than a selection based on a single attribute. The efficacy of our method was demonstrated in three patients by showing that the top predicted tumor-specific clonotypes were tumor-reactive. The fact that not all initially predicted tumor-specific TILs (patients 1,2, **Fig. 1B,C right**) were expanded and showed responses to tumor challenge can probably directly be attributed to exhaustion of the T cells[36]. In patient 3, TCR-T cells generated with TCRs of the top four clonotypes proved to be tumor-specific and three of them recognized a neoepitope derived from oncoprotein KRAS Q61H. Predicted tumor-specific clonotypes from five additional patients showed congruent differentiation trajectories as determined by scRNA-Seq.

Regarding the clinical relevance of the KRAS-epitope, driver mutations in TP53, EGFR, and KRAS invariably represent clonal (or truncal) mutations in smoking- and non-smoking-related lung cancer[51]. Resultant expression in all tumor cells in combination with the high immunogenicity of the epitope make the KRAS Q61H epitope an attractive target for immunotherapy. Immunogenicity is inferred by the fact that homologous NRAS mutations (ILDTAG**K**EEY, ILDTAG**R**EEY) have been shown to be immunogenic in HLA-A*01:01-positive melanoma patients and presentation of the peptides was shown by immunopeptidomics[52]. In our study, three independent T-cell clonotypes with strong avidity targeting epitope ILDTAG**H**EEY were found in the patient and highly homologous TCRs were discovered infiltrating KRAS Q61H positive tumors in other patients. Moreover, all TCRs were rediscovered in a relapse lesion and one even in circulating cell-free DNA analyzed almost three years after surgery of the primary tumor. The antigenic peptide is presented by HLA-A*01:01, which is expressed in 23,7% of tumors in the TCGA database[25]. Compared with other KRAS-driver mutations, such as G12-hotspot mutations, the Q61H mutation is rare (according to TCGA occurring in lung, colorectal, and pancreatic cancer in 0.2%, 0.7%, 2.8% of cases, respectively), which may explain why this immunogenic epitope has remained undetected so far[53]. However, given the high incidences of the mentioned tumor indications in Europe and the United States, hundreds of patients per year would be eligible for ACT with TCR-T cells expressing these TCRs. Clinical responses comparable to those reported for a small number of patients using TCR-T cells transduced with KRAS G12-mutation specific TCRs can be expected[29, 31, 53].

In conclusion, discovery of the tumor-specific and mutant KRAS-reactive TCRs in the presented cases implies that our strategy to identify and select tumor-specific TCRs can be applied to many patients with different tumors, provided that surgical material for analysis is available. Synthesis and cloning of the natural TCRs and manufacturing of autologous T cells with these TCRs for therapeutic application can be expected to be safe because the tumor-reactive clonotypes have passed thymic selection and dealt with the tumor in vivo without apparent adverse effects. Moreover, ACT with T cells transduced with three to four dominant tumor-specific TCRs per patient can address tumor heterogeneity and counteract immune-escape mechanisms[54, 55]. However, while developing such personalized TCR-T cell products is feasible, the clinical implementation is challenging from a manufacturing and regulatory perspective[53]. Yet, to overcome the challenges is worthwhile because an approach for the direct selection of tumor-specific TCRs from TILs can make more patients with solid tumors eligible for TCR-T cell therapy than antigen-centered selection approaches.

## Acknowledgments

We are grateful to Dr. Patricia Haehnel and Dr. Sigrid Horn (University Medical Clinic UMC, Mainz, Germany) for providing us with the NSCLC cell lines NCI-H460 and MZ-LC-16, respectively. Buffy coat preparations from blood of healthy donors were provided by the blood bank of UMC Mainz. Part of the present work is a component of CD’s doctoral thesis. Throughout the experimental part of his thesis, CD received scientific advice and training by the Mainz Research School of Translational Biomedicine (TransMed, https://www.unimedizin-mainz.de/transmed/home.html). Finally, we are grateful to donors and patients who contributed materials critical to the accomplishment of this work.

## Author contributions

Conceptualization: VL, CD, SH, TW, VS, RH, DW

Surgery and Pathology: SE, CB, HH, KK

Methodology: CD, MF, AD, SS, CW, NG, SP, KG, DB

Investigation: KG, DB, VL, SH, HH, RH, PA, SWL, DW

Visualization: VL, CD, HH, KG, DB, SH

Project administration: SH, PA, DW, TW

Supervision: VL, CW, TW, SH, DW, PA, RH

Writing – original draft: VL, CD, SH

Writing – review & editing: VL, CD, SH, RH, VS, PA, TW

## Declaration of interests

SH, VS and RH are co-founders and shareholders of HS Diagnomics GmbH and TheryCell GmbH and have filed patent applications for technologies applied in the study (EP2746405B1, EP3180433B1). SH serves as the CEO and VL as the CSO of HS Diagnomics and Therycell and AD, SS, and VL are shareholders of both companies. All other authors have declared that no conflicts of interest exist.

## Materials and Methods

### Key Resources Table

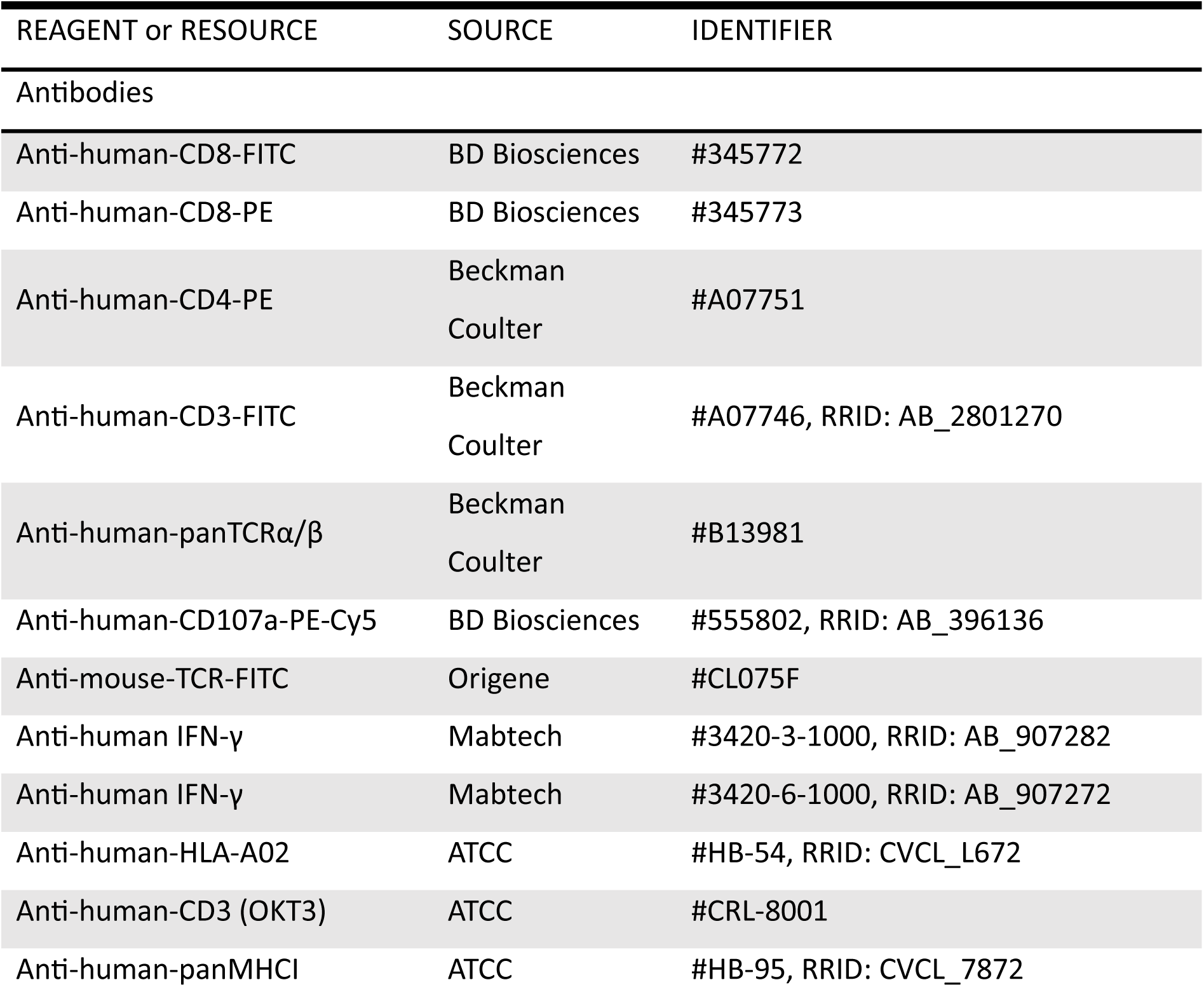
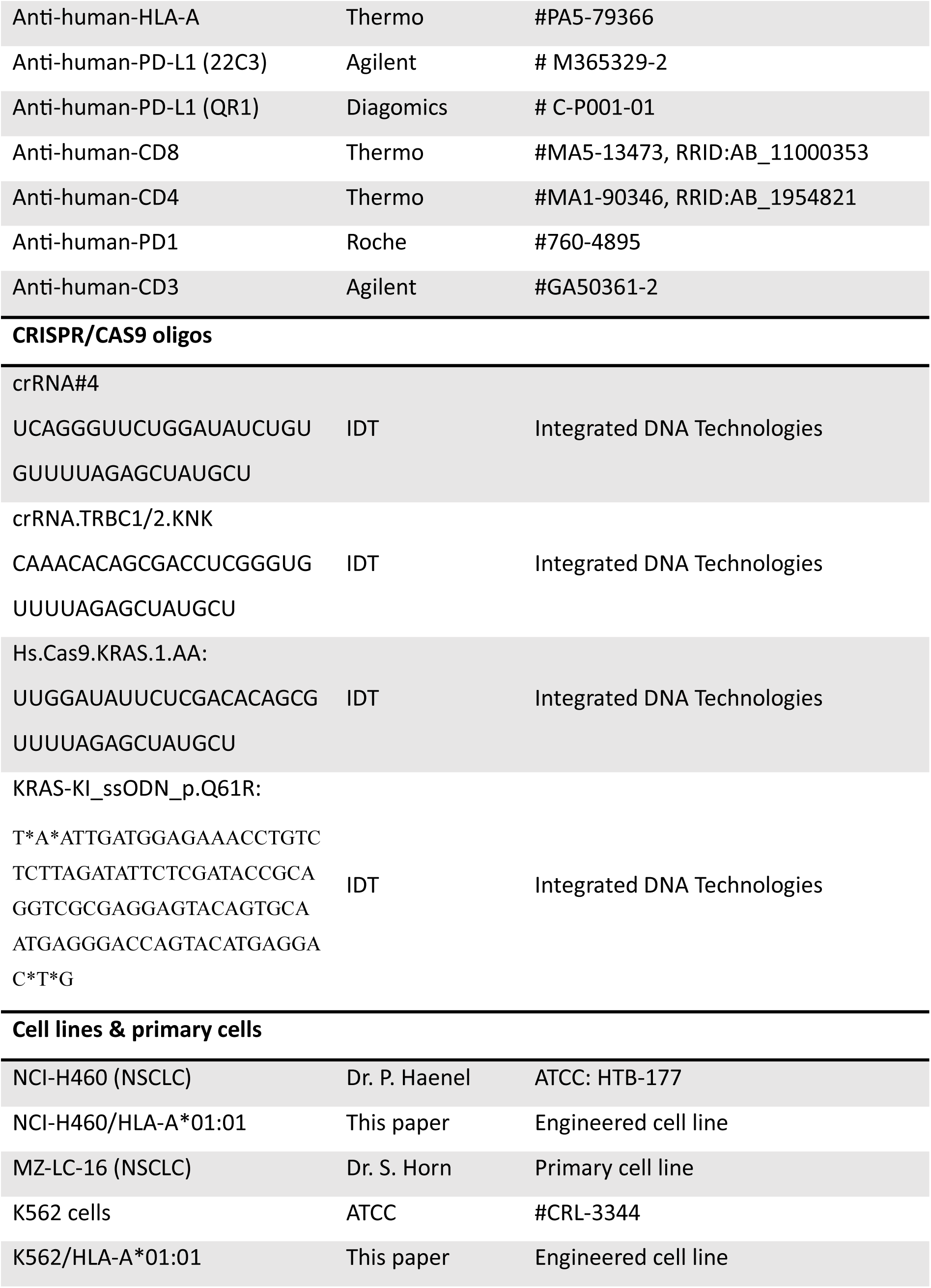
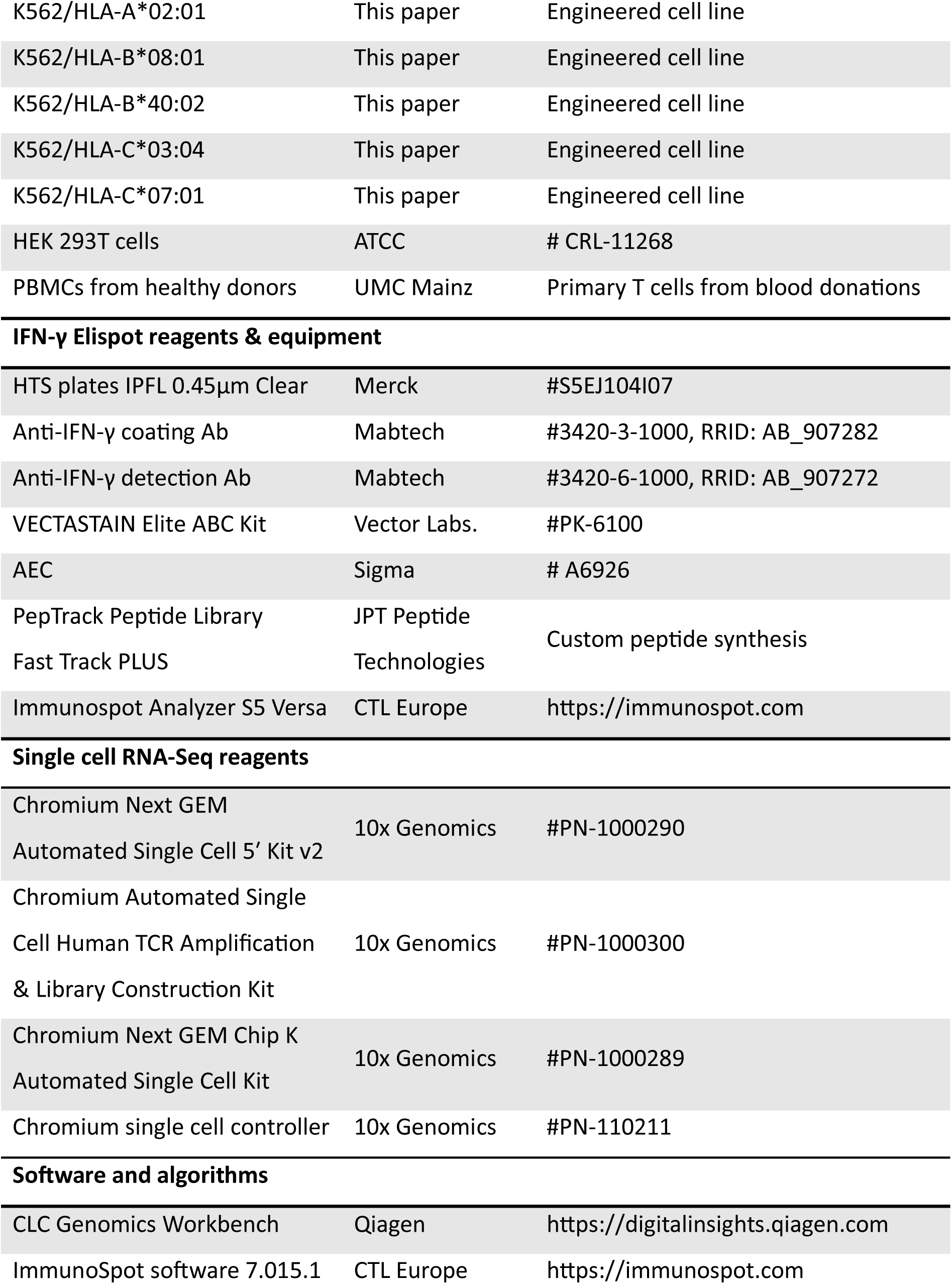
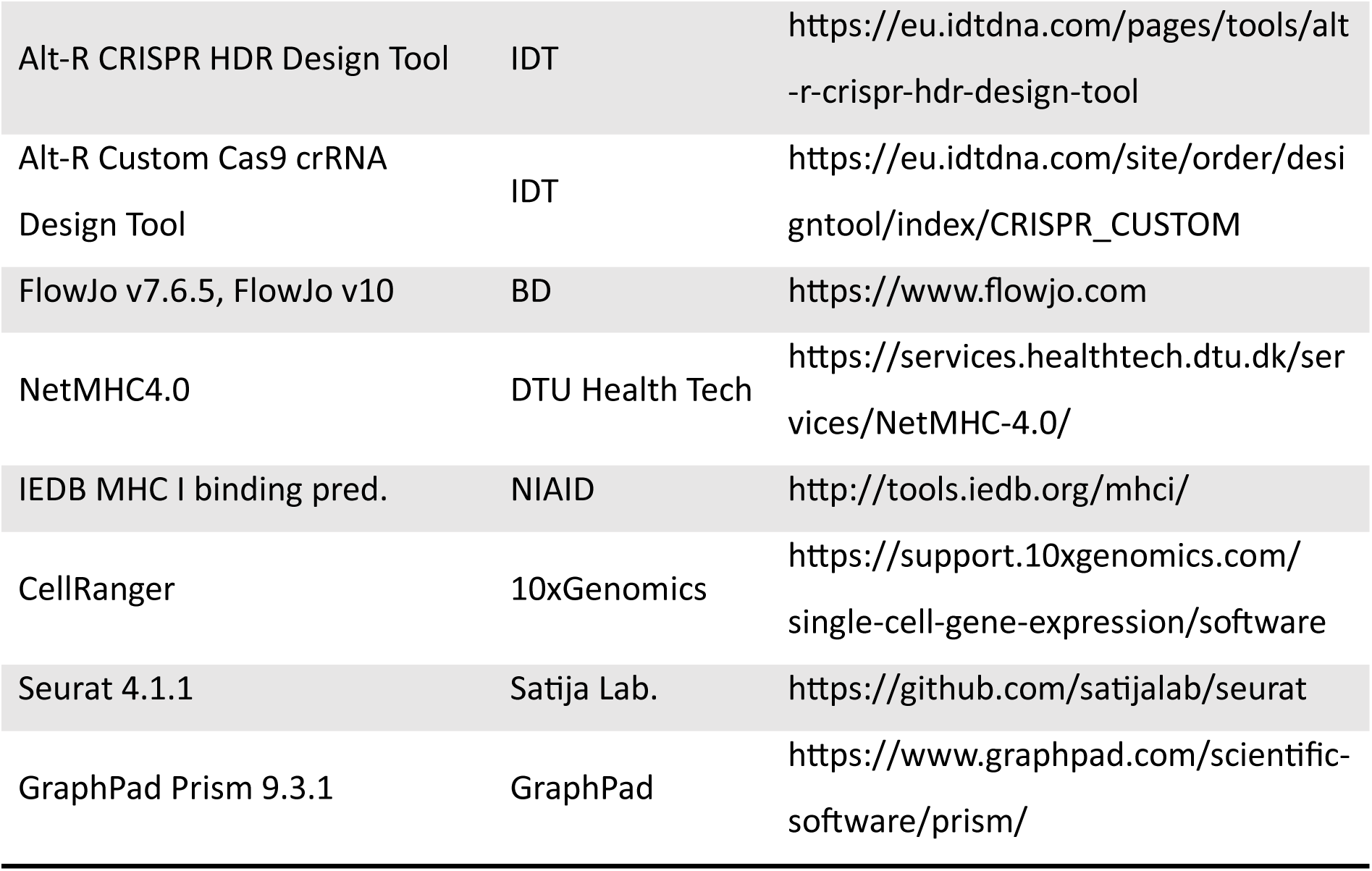

### Resources Availability

#### Lead contact

Further information and requests for resources and reagents should be directed to and will be fulfilled by lead contact Volker Lennerz.

#### Materials availability

This study did not generate new unique reagents.

#### Data and code availability

Any additional data required to reanalyze the data reported in this work is available from the lead contact upon request.

#### Patient material

From seven patients with NSCLC, clinical material including blood, fresh tumor- and adjacent normal-lung tissue selected by a pathologist was obtained and transported to the lab in the fastest way possible. Lymphocyte subpopulations (CD3, CD4, CD8, PD1) were isolated from fresh tumor, lung, and blood for high throughput TCR-VDJ amplification and TCR-repertoire profiling (TCRSeq). From six patients, including patients 2 and 3, part of the CD3 fractions underwent single cell RNA-Seq (see below). For patients 1, 2, and 3 functional assays were performed. Only in patient 3, in addition to samples from primary surgery, FFPE samples from relapse surgery in January 2021 as well as blood samples collected in September and December 2021 were investigated. The clinical course of patient 3 and derivation of all samples are summarized in **Fig. S1**. Similarly, patient 3 was the only subject in whom lung and tumor tissue samples were subjected to whole exome and transcriptome sequencing, and cell-free (cf)DNA was isolated from blood plasma and also subjected to TCR repertoire profiling.

The study was performed in accordance with the declaration of Helsinki. Sample acquisition from NSCLC patients was approved by the ethics committee of the Aerztekammer Berlin (Eth-08/18) and informed consent received from all patients.

#### Primary tissue and blood cell processing

Small pieces from different tissue regions (∼1g) each of tumor and adjacent normal tissue were physically disrupted using scalpels and subjected to GentleMACS tissue dissociation according to protocol (Miltenyi Biotec, Bergisch-Gladbach, Germany). After filtering through a 70 μm-cell strainer, one aliquot each of the cell suspension underwent Percoll gradient centrifugation and the remainder was cryopreserved. Percoll-interphases were collected and rested overnight at 0.5Ö10^6^ cells/ml in TexMACS medium (Miltenyi Biotec) plus 25 mM HEPES (pH 7.2), L-glutamine (Lonza, Köln, Germany), 50 mM beta-mercaptoethanol (ThermoFisher Scientific, Waltham, MA, USA), and 10% autologous serum. Tumor- and lung cell pellets were resuspended and cryopreserved. After harvesting and washing the leukocyte fractions, CD3-, CD4-, CD8-, and PD1-positive cells were isolated from TILs and lung-leukocytes using magnetic beads (Miltenyi Biotec) or FACS. For whole exome sequencing and RNA-Seq (see below), sections of different tumor and lung regions were pooled separately, and snap-frozen in liquid nitrogen until preparation of nucleic acids.

#### TCR repertoire profiling of tumor- and lung-infiltrating lymphocytes and TIL-scRNA-Seq

From sorted subpopulations of TILs, lung-infiltrating lymphocytes, PBMCs, and blood plasma, genomic (g)DNA was isolated and subjected to TCR-VDJ-amplification using human TRBV/J-specific primer sets and NGS-analysis (referred to as TCRseq) [44, 45]. Briefly, gDNA from CD3-, CD8-, and PD1-positive T-cell subpopulations was isolated using the QIAamp blood kit (Qiagen, Hilden Germany) and NGS libraries were generated employing a two-step PCR protocol [44]. gDNA from FFPE samples processed with the AllPrep DNA/RNA FFPE kit (Qiagen) and from urine and plasma with the Norgen Plasma/Serum RNA/DNA Purification Mini Kit (BioCat GmbH, Heidelberg, Germany) was applied to TCRseq, too. In addition to TRBV-sequencing, FFPE-samples from 29 patients with various tumors expressing KRAS Q61H were subjected also to TCRseq using human TRAV/J-specific primer sets. Single cell cDNA-libraries were generated from CD3+ TIL single cell suspensions using 10x Genomics® GemCodeTM Technology (10x Genomics B.V., Leiden, The Netherlands). Briefly, lymphocytes were processed using the 10x Genomics Chromium Next GEM Single Cell V(D)J Reagent Kit in combination with the Chromium Single Cell V(D)J Enrichment Kit (Human) according to protocol. After clean-up, libraries were analyzed by Illumina next generation sequencing (StarSEQ GmbH, Mainz, Germany).

#### CRISPR/CAS9 engineering of primary T cells and cell lines

T cells isolated from Buffy Coats from three healthy donors were isolated by Ficoll (Sigma-Aldrich, Taufkirchen, Germany) separation and MACS-sorting according to protocol (Miltenyi Biotec). After OKT3-activation (plate-bound, 30 ng/µl), T cells were subjected to CRISPR/CAS9-mediated knockout of endogenous TCRs. Ribonucleoprotein (RNP) complexes were delivered by Human T Cell Nucleofector^TM^ Kit (Lonza, Basel, Switzerland). Both TCR chains were targeted by two crRNAs. The TRBC-crRNA was previously described [56], the TRAC-crRNA was designed using the Alt-R Custom Cas9 crRNA Design Tool (IDT, Coralville, USA). Combined at 1:1-ratio with the Alt-R^®^ CRISPR-Cas9 tracrRNA (IDT) two different gRNA complexes were formed. gRNA complexes were combined with recombinant Cas9 (IDT) for RNP complex generation (20 min, RT). 4Ö10^6^ T cells were transfected in Nucleofector^®^ Solution supplemented with 1 µM Alt-R^®^ Cas9 electroporation enhancer (IDT) and 4 µM of RNPs. T cells were electroporated with program T-023 on a Nucleofector^TM^ 2b Device (Lonza). T cells were cultivated at 1Ö10^6^ cells/ml Panserin complete (plus 600 U/ml IL-2). TCR-KO efficiency was assessed 4-6 days later via flow cytometry. CRISPR/CAS9 genome editing was also used to substitute KRAS-codon 181-183 CGC (encoding KRAS_p.Q61R) for KRAS-codon 181-183 CAT (encoding KRAS_p.Q61H) in HLA-A*01:01-expressing NCI-H460 NSCLC cells by homology-directed repair (HDR). A specific crRNA was designed by the Alt-R Custom Cas9 crRNA Design Tool (IDT) and then combined with Alt-R^®^ CRISPR-Cas9 tracrRNA (IDT) to generate specific RNPs. ssODN constructs encoding the KRAS_p.Q61R mutation flanked by homology arms of 40-46 nt were generated. ssODN constructs were stabilized by IDT-proprietary end-blocking groups and two phosphorothioate bonds. Three silent mutations at ssODN positions 48, 60 and 63 prevented the Cas9 enzyme from re-cutting target sequences after HDR. Cells underwent nucleofection with RNP complexes using program X-001 (Lonza). Briefly, two-part gRNA complexes were prepared at 100 µM and combined with recombinant Cas9-NLS nuclease (QB_3_ Macrolab, Berkeley, USA) for the generation of KRAS-specific RNPs. After formation (20 min, RT), 3Ö10^6^ NCI-H460/HLA-A1 cells were transfected in 110 µl OptiMEM supplemented with 4 µM ssODN templates, 1 µM Alt-R^®^ Cas9 enhancer (IDT) and 4 µM RNPs. Nucleofected cells were cultivated in 2 ml RPMI supplemented with 30 µM Alt-R^TM^ HDR Enhancer V2 (IDT) per well of a 6-well plate. Nucleofection medium was replaced by RPMI+ after 18-20 h. Clonal cell lines were established via limiting dilution cloning.

#### TCR-encoding DNA-synthesis and cloning

T-cell receptor alpha-chain (TRAV/J-) and TCR beta-chain (TRBV/D/J-) region-coding sequences were synthesized as G-blocks (IDT) and cloned as bicistronic constructs connected by a P2A-encoding linker into gamma-retroviral expression vector pMX-puro as described [57]. TCRs were designed as chimeric constructs in which the human TRA- and TRB-constant-region sequences (TRAC, TRBC) were replaced by murine homologous sequences.

#### Stable transduction of primary T cells and cell lines

TCR-encoding γ-retroviral particles were produced for transduction of primary T cells as described [47, 57]. Briefly, T cells from Buffy coats of healthy donors were isolated by Ficoll separation followed by CD8- and CD4-magnetic bead isolation according to protocol (Miltenyi Biotec). After activation with plate-bound OKT-3 (30ng/μl) and culture for 3-5 days, endogenous TRAC/TRBC-knockout was accomplished. Viral particles were produced using Phoenix-ampho packaging cells seeded at 1,3×10e6 cells per 100mm plate. After 24 hours, cells were co-transfected with 5μg each pCOLT-GALV, pHIT60 and 10μg pMX/TCR using Fugene-6 according to protocol (Promega, Madison, WI, USA). Transfection medium was changed for T-cell medium after 24h and supernatant was harvested 16 hours later following cell-pelleting. TCR-T cells were generated by spin-inoculation with retrovirus-containing T cell medium of 2×10e6 T cells per reaction and TCR-T cells expanded and selected using puromycin (1μg/ml, Sigma Aldrich) as reported [47, 57]. Transductions of K562- and NCI-H460 cells were performed accordingly with pMX/HLA-constructs.

#### Computational analyses

The results of TCR repertoire sequencing (TRB or TRA chains, 2x 150bp paired Illumina reads) were processed by in-house developed software using both reads to build a consensus sequence covering the complete CDR3 region and removing inconsistent non-overlapping read-pairs. High-quality consensus CDR3 sequences were clustered into unique clonotypes, respective V- and J-segment IDs, and a clonotype frequency was calculated as percentage of clonotype reads compared to all sample reads. Clonotype sequences were further analyzed for productive ORFs discarding non-functional sequences. The output frequency matrix with each row belonging to a unique CDR3 nucleotide sequence (**Table S4**) showed clonotype frequencies in peripheral blood, tumor and adjacent non-tumor tissues. A frequency ratio TIL/adjacent normal lung was calculated for all clonotypes; those with a ratio >5 were enriched for tumor-specific T-cells[58].

Single cell sequencing reads were processed with the 10X Genomics Cell Ranger pipeline (v6.0.2) with default parameters to demultiplex and generate unique molecular identifier (UMI) matrices. The matrices were used in R with Seurat (v4.1.1) for quality control and downstream analyses. Each sample matrix was individually inspected for quality control before integration into a merged dataset. Cells with less than 400 UMI, fewer than 250 genes and greater than 20% UMI in mitochondrial genes were removed. For sc-gene expression analysis, TCR genes were neglected to avoid clustering based on certain V or J gene segments. To account for library chemistry and align cells from different samples, an integration method based on highly variable shared genes was used. Starting with the SCTransform function for normalization and identification of the most variable genes, we also regressed out variation due to mitochondrial expression. The top 3000 variable genes were used from the SCTransform object to find “anchors” with the FindIntegrationAnchors function and thereafter processed with IntegrateData to produce a sample-corrected data set. The first 30 principal components of the integrated data were used for uniform manifold approximation and projection (UMAP) construction, as well as the unsupervised graph-based clustering to identify distinct groups of cells, including CD4- and CD8-positive T-cell clusters. A subclustering of CD8-associated clusters (CD8-GZMK, CD8-ZNF683, CD8-ENTPD1 and CD4/8-MT) was done with the same parameters.

#### Culture of cell lines, TILs and primary T cells

Cell lines and primary T cells were grown in incubators at 37°C, 5% CO_2_, >85% humidity. HEK 293T-, K562-, NCI-H460-, and MZ-LC-16 cells (kindly provided by Dr. P. Haenel and Dr. S. Horn, UMC, Mainz, Germany) were maintained in RPMI-1640 supplemented with 10% FBS, and 1% penicillin/streptomycin (RPMI+, Sigma Aldrich, Taufkirchen, Germany). HLA-monoallelic K562 cells were engineered to express all six HLA-alleles of patient 3, and NCI-H460 to express HLA-A*01:01 using γ-retroviral transduction. Cells were maintained in RPMI+ plus puromycin (1μg/ml, Sigma Aldrich). Primary T cells from healthy donors were grown in Panserin-413 (PAN-Biotech, Aidenbach, Germany) supplemented with 10% heat-inactivated pooled human serum (provided by the blood bank of UMC Mainz, Germany), 1% Penicillin/Streptomycin (Sigma Aldrich), and rhIL-2 (250-600 IU/mL Novartis, Basel, Switzerland). Cell lines underwent STR-analysis for identity verification and were regularly subjected to mycoplasma testing to ensure absence of contamination. In patients 1 and 2, parts of the over-night rested TILs were taken in culture and expanded for two to three weeks. The procedures are outlined in Figures **Fig. S3** (patient 1) and **Fig. S4** (patient 2) and further details are provided in the legends.

#### Flow cytometry

T lymphocyte subpopulations were stained with monoclonal antibodies anti-CD8-FITC or -PE (clone 9.11, SK1; BD Biosciences, Heidelberg, Germany), anti-CD4-PE (13B8.2), anti-CD3 (UCHT1), anti-human TCR constant domain (IP26A; all Beckman Coulter, Krefeld, Germany), and anti-murine TCR constant domain (FITC, CL075F, Origene, Rockville, MD, USA). TCR-T cells were tested for activation-induced cytolytic responses by coincubation with target cells (1:1) overnight followed by staining with anti-murine TCR constant domain antibody (FITC, CL075F, Origene) and anti-CD107a mAb (PE-Cy5, clone H4A3, BD Biosciences, Heidelberg, Germany). Antibody-stained cells were analyzed on either FACS Canto II or Melody instruments (BD Biosciences). Data were analyzed using FlowJo 10 analysis software (BD Biosciences). Additional antibodies used in the study are listed in the Key Resources Table.

#### Whole exome- and RNA-sequencing of lung- and tumor tissue nucleic acid preparations

Genomic DNA for WES and totalRNA for RNA-Seq were isolated from frozen tumor and lung tissues using the QIAamp Fast DNA Tissue Kit and RNeasy Plus Kit as per protocols (Qiagen). Briefly, tissue blocks were cut into chunks of approx. 25 mg by using pre-chilled mortars and RNase-free scalpels. To prevent degradation by RNases, samples were kept cold with liquid nitrogen and mortars were sterilized beforehand by baking for 6 h at 180°C. To address tumor heterogeneity, three individual lung and tumor tissue chunks obtained from different areas were used for every gDNA and RNA purification. Tissue fractions were homogenized in QIAzol Lysis Reagent (Qiagen) using a TissueLyser LT bead mill (Qiagen) according to protocol (50 Hz, 2x 2.5 min). Exome enrichment, preparation of sequencing libraries and NGS were with StarSEQ GmbH (Mainz, Germany). Raw data were processed and analyzed using CLC Genomics Workbench (https://digitalinsights.qiagen.com/).

#### Transient transfection of HEK 293T cells

Overexpressed antigen- and KRAS-encoding cDNAs were reverse-transcribed and amplified from patient 3 tumor- and NCI-H460 cell RNAs using standard RT- and PCR-kits and cloned into pcDNA3.1-derived vectors employing Gateway technology (Invitrogen). HEK 293T cells were transiently transfected with plasmids encoding HLA-A*01:01 and wildtype and mutated KRAS full-length and fragment cDNAs using Lipofectamine 2000 as per protocol (Invitrogen). Briefly, transfection was carried out in wells of Multiscreen HTS 96-well plates previously prepared for ELISpot testing. Per well, 20,000 cells were transfected with 100ng HLA-plasmid, 300ng antigen-encoding plasmid and 0.5μl Lipofectamine transfection reagent. Twenty-four hours after transfection, recombinant 293T cells were used as target cells in IFN-γ ELISpot assays.

#### IFN-γ ELISpot assays

Response analyses of TCR-T cells by IFN-γ ELISpot assays were performed as reported [59]. TCR-T cells were expanded for several weeks and aliquots cryopreserved every week. TCR-T cells were ready for testing when they exhibited >50% cTCR-expression. ELISpot assays were performed with TCR-T cells from culture or after thawing. Thawed cells were rested overnight before testing. Briefly, HLA- and antigen-cDNA transfected 293T cells (20000/well), peptide pulsed (2μM) K562/HLA (50,000/well), NCI-H460, NCI-H460/HLA-A*01:01 cells (50,000/well), or tumor- and lung cell suspensions (20,000/well) were co-incubated with TCR-T cells (2,000-10,000 cTCR-positive cells/well) in IFN-γ antibody-coated Multiscreen HTS plates (Merck-Millipore, Darmstadt, Germany) overnight (16-20h). Positive control OKT3-antibody (purified from hybridoma, 400ng/ml) was co-coated in control-wells together with the anti-IFN-γ-antibody. All reactions were set-up in duplicates or triplicates. After 20h, cells were discarded, and tests developed as per protocol [59]. Plates were scanned and analyzed by ImmunoSpot Analyzer S5 Versa with ImmunoSpot software 7.0.15.1 (CTL Europe, Bonn, Germany). For peptide-pulsing of K562/HLA-monoallelic cell lines, a Fast Track Peptide Library of candidate neoepitopes was purchased from JPT Peptide Technologies (Berlin, Germany). Lyophilized peptide pools were reconstituted with DMSO and after dilution with RPMI (16μg/ml RPMI/5% DMSO) stored at −20°C. Pan-HLA antibody W6/32 (purified from hybridoma supernatants) was used to block pMHC-specific recognition of target cells by TCR-T cells.

#### Immunohistochemistry

Primary tumor- and relapse FFPE samples were analyzed for tumor areas (H&E staining) and tumor cell expression of HLA-A using a polyclonal antibody against a common epitope of all HLA-A alleles (Thermo Fisher Scientific, Dreieich, Germany) and of PD-L1 using monoclonal (m)Abs 22C3 (Agilent, Waldbronn, Germany) and QR1 (Diagomics, Cedex, France). On consecutive slices, T cells were stained with mAbs specific for CD8 (C8/114B, Thermo Fisher Scientific), CD4 (4B12, Thermo Fisher Scientific), and PD-1 (NAT105, Roche, Rotkreuz, Switzerland). A polyclonal Ab was used to detect CD3 (DAKO, Agilent, Waldbronn, Germany). For identifiers of all IHC-antibodies used see key resources table.

## List of Supplementary Tables

Table S1 (Clinical data patient 3)

Table S2 (Experimentally confirmed tumor-specific T-cell clonotypes of patient 1)

Table S2 (Experimentally confirmed tumor-specific T-cell clonotypes of patient 2)

Table S4 (TCRseq results patient 3)

Table S5 (10X Genomics scTCRseq results patient 3)

Table S6 (NGS results WES and scRNA-Seq of patient 3, SNVs, InDels)

Table S7 (Peptide binding predictions of neoantigens discovered in patient 3’s tumor)

Table S8 (Homologous TCR-CDR3 sequences to KRAS Q61H-TCRs in 29 KRAS Q61H-positive tumors, Fig. 5B)

Table S9 (Resource table of raw data for the generation of figures)

