## Supplementary material for "T-cell receptors identified by a personalized antigen-agnostic screening approach target shared neoantigen KRAS Q61H": Table S3

| **Score** | **TRB-CDR3 peptide** | **Freq. tumor** | **Freq. lung** | **Tumor / lung** | **Freq. PD1+** | **Selected** | **Tumor-reactive^1^** |
| --- | --- | --- | --- | --- | --- | --- | --- |
| 1 | CASSLGTAGEQFF | 6.310 | 0.296 | 21.35 | 3.520 | + | + (3.443) |
| 2 | CAISGLLDAGDPDTQYF | 2.990 | 0.052 | 57.29 | 1.743 | + | + (0.215) |
| 3 | CASSQEPVSSYNSPLHF | 2.393 | 0.128 | 18.75 | 1.001 | + | + (3.431) |
| 4 | CASSRTGFEANTEAFF | 2.011 | 1.040 | 1.93 | 1.248 | - | 0.000 |
| 5 | CASSQGDRGRENSPLHF | 1.882 | 0.602 | 3.12 | 0.446 | - | + (21.431) |
| 6 | CASSPGQGDYEQYF | 1.796 | 0.063 | 28.46 | 1.055 | + | 0.000 |
| 7 | CASSLWTSADTQYF | 1.796 | 29.810 | 0.06 | 0.151 | - | + (5.235) |
| 8 | CAWSGGQGRLNQPQHF | 1.639 | 0.205 | 8.00 | 0.193 | + | + (0.390) |
| 9 | CASSALGDTEAFF | 1.371 | 0.000 | >1000 | 0.144 | + | 0.000 |
| 10 | CASSIIGTGDQYF | 1.314 | 0.086 | 15.29 | 0.258 | + | 0.000 |
| 11 | CAISTVEADTIYF | 1.309 | 0.005 | 276.10 | 0.640 | + | 0.000 |
| 12 | CASSYLTDTQYF | 1.172 | 0.748 | 1.56 | 0.731 | - | 0.000 |
| 13 | CASSPWTDGGYTF | 1.104 | 0.253 | 4.36 | 0.052 | - | 0.000 |
| 14 | CASSWGTTSPLHF | 1.060 | 0.049 | 21.48 | 0.241 | + | 0.000 |
| 15 | CASSPTTGDEQYF | 1.015 | 0.042 | 24.32 | 0.835 | + | 0.000 |
| 16 | CAWEPGHLGETQYF | 0.981 | 0.050 | 19.68 | 0.056 | - | 0.079 |
| 17 | CASSPVGGYTEAFF | 0.979 | 0.572 | 1.71 | 0.054 | - | 0.000 |
| 18 | CASSIGKTAYEQYF | 0.776 | 0.019 | 40.90 | 0.000 | - | + (2.639) |
| 19 | CASSLGPAGNTIYF | 0.758 | 0.106 | 7.12 | 0.111 | - | + (3.547) |
| 20 | CASSSLGQPNTEAFF | 0.663 | 0.003 | 199.83 | 0.401 | - | 0.000 |

^1^ CD137-positive sorted cells after stimulation of expanded TILs with autologous tumor cells. Positivity threshold: 0.1%.

Table S3: Top 20 TIL clonotypes of patient 2: Frequencies (percentages) of the top 20 TILs in comparison to additional fractions of the primary tumor and after expansion and tumor challenge.
