## Supplementary material for "T-cell receptors identified by a personalized antigen-agnostic screening approach target shared neoantigen KRAS Q61H": Table S2

| **Score** | **TRB-CDR3 peptide** | **Freq. tumor** | **Freq. lung** | **Tumor / lung** | **Freq. PD1+** | **Selected** | **Tumor-reactive^1^** |
| --- | --- | --- | --- | --- | --- | --- | --- |
| 1 | CASSVDRGAEAFF | 3.6602 | 0.4070 | 8.9903 | 2.0895 | + | + (2.7417) |
| 2 | CAWNKQVDGYTF | 2.3939 | 0.0546 | 43.7316 | 0.8854 | + | + (0.7399) |
| 3 | CASSFGVMNTEAFF | 2.3298 | 1.4606 | 1.5951 | 0.5585 |  | 0.0893 |
| 4 | CASSPDGETQYF | 2.2556 | 0.2012 | 11.2029 | 0.8427 | + | + (1.1265) |
| 5 | CASSLGQAYEQYF | 1.7864 | 1.0162 | 1.7578 | 0.1642 |  | 0.0449 |
| 6 | CASSPVAGMNTEAFF | 1.4172 | 3.3855 | 0.4186 | 0.2980 |  | + (0.2441) |
| 7 | CAISDWTGSNYGYTF | 1.1625 | 0.0583 | 19.9125 | 0.2638 | + | + (2.3817) |
| 8 | CASSGRGDLLEQYF | 1.1208 | 0.5964 | 1.8788 | 0.0858 |  | 0.0000 |
| 9 | CASSETGAAETQYF | 1.0946 | 4.8834 | 0.2241 | 0.2134 |  | 0.0971 |
| 10 | CASSRLAGGTDTQYF | 0.9505 | 2.4768 | 0.3837 | 0.0923 |  | 0.0399 |
| 11 | CASSSGLVYEQYF | 0.8981 | 0.1284 | 6.9893 | 0.4463 | + | + (0.8880) |
| 12 | CASSTGTGGLGELFF | 0.8748 | 0.1020 | 8.5688 | 0.7057 | + | + (0.6506) |
| 13 | CASSEAPPLYYEQYF | 0.8549 | 0.2113 | 4.0447 | 0.0000 |  | 0.0000 |
| 14 | CASSNDRAGLNEQFF | 0.8461 | 0.7394 | 1.1442 | 0.0240 |  | 0.0000 |
| 15 | CATSDGRLEQFF | 0.8219 | 0.1129 | 7.2728 | 0.8480 | + | 0.0000 |
| 16 | CASSLGYRYGTEAFF | 0.8103 | 3.5121 | 0.2307 | 0.2943 |  | 0.0000 |
| 17 | CASSQDNGGYGYTF | 0.8030 | 0.1484 | 5.4062 | 0.1943 | + | 0.0000 |
| 18 | CASSQGDSFYGYTF | 0.8001 | 0.1949 | 4.1037 | 0.0394 |  | 0.0000 |
| 19 | CASSADLGDRVNGYTF | 0.7821 | 0.8068 | 0.9693 | 0.2459 |  | 0.0000 |
| 20 | CASSLDRGGYEQYF | 0.7540 | 0.1402 | 5.3728 | 0.1224 |  | 0.0000 |

^1^ IFN-γ-positive captured cells after stimulation of expanded TILs with autologous tumor cells. Positivity threshold: 0.1%.

Table S2: Top 20 TIL clonotypes of patient 1: Frequencies (percentages) of the top 20 TILs in comparison to additional fractions of the primary tumor and after expansion and tumor challenge.
