## Supplementary material for "T-cell receptors identified by a personalized antigen-agnostic screening approach target shared neoantigen KRAS Q61H": Table S1

| Time  (mo, yr) | Diagnosis,  clinical findings | Treatment | Staging information | Molecular Pathology |
| --- | --- | --- | --- | --- |
| Jun 2018 | lung adeno-carcinoma,  left upper lobe | lobectomy left upper lobe and mediastinal lymph node dissection | pT2a pN0 (0/24) pL1 pV1 R0 cM0;  UICC IB | PD-L1 55% (TPS) |
| Jul-Sep 2019 | local recurrence | chemoradiotherapy (carboplatin, vinorelbine; total irradiation dose 66 Gy) | - | - |
| Oct 2019-Nov 2020 | - | durvalumab maintenance | - | - |
| Jan 2021 | progressive  local recurrence,  no evidence of metastatic disease | extended left pneumonectomy | yrpT2a yrpN0 (0/11);  regression grade IIa (≥ 10% vital tumor tissue)* | *KRAS* c.183A>T/p.Q61H;  *TP53* c.638G>A/p.R213Q;  *SMO* c.1225G>T/p.G409C;  PD-L1 50-60% (TPS) |
| Since 2021 until now | No evidence of disease |  |  |  |

**T**a**ble S1: Clinical course of patient 3**

*: according to Junker et al., doi: 10.1378/chest.120.5.1584
