## Supplementary information for "T-cell receptors identified by a personalized antigen-agnostic screening approach target shared neoantigen KRAS Q61H"

### Slide 1
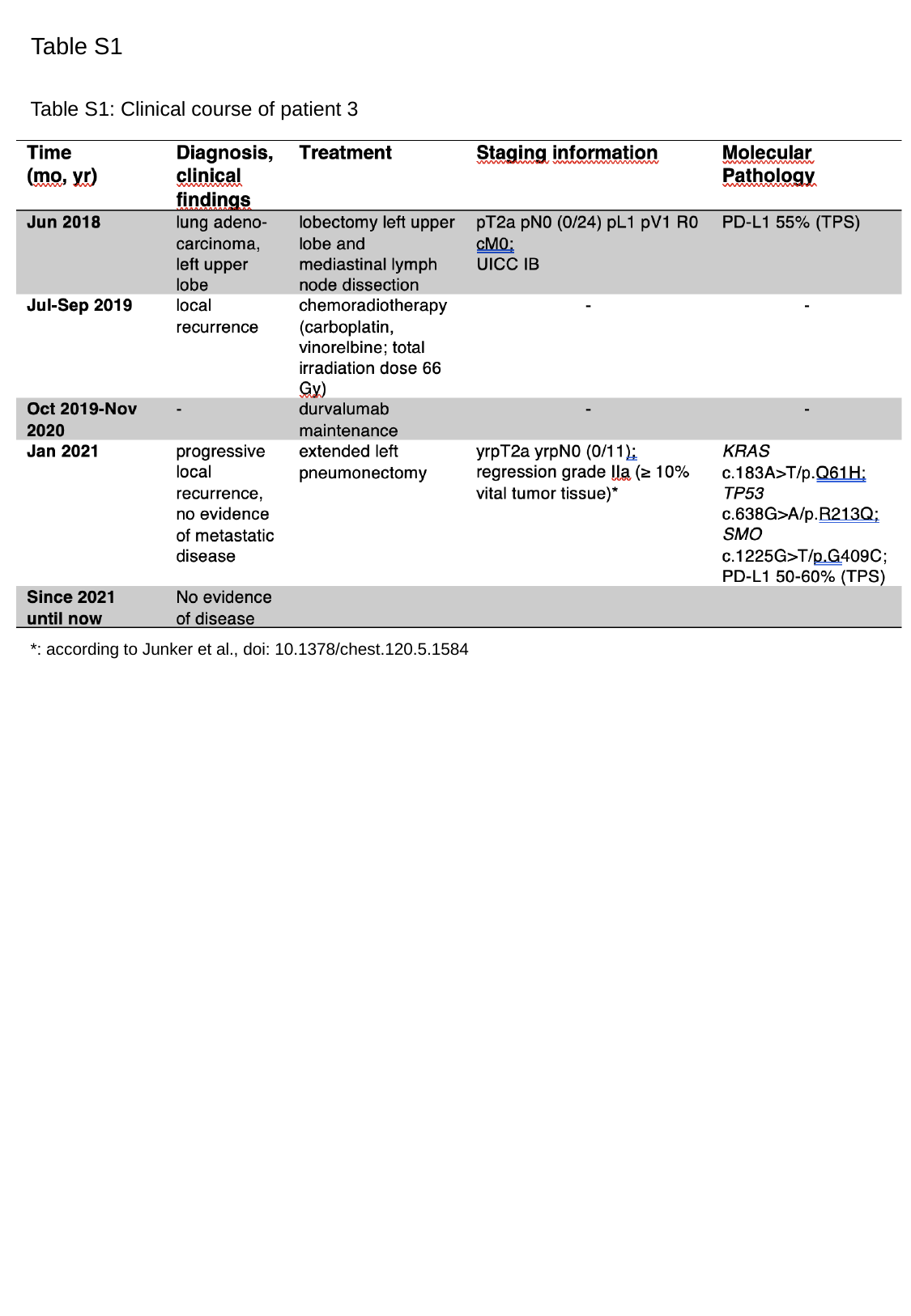

Table S1
Table S1: Clinical course of patient 3
*: according to Junker et al., doi: 10.1378/chest.120.5.1584

### Slide 2
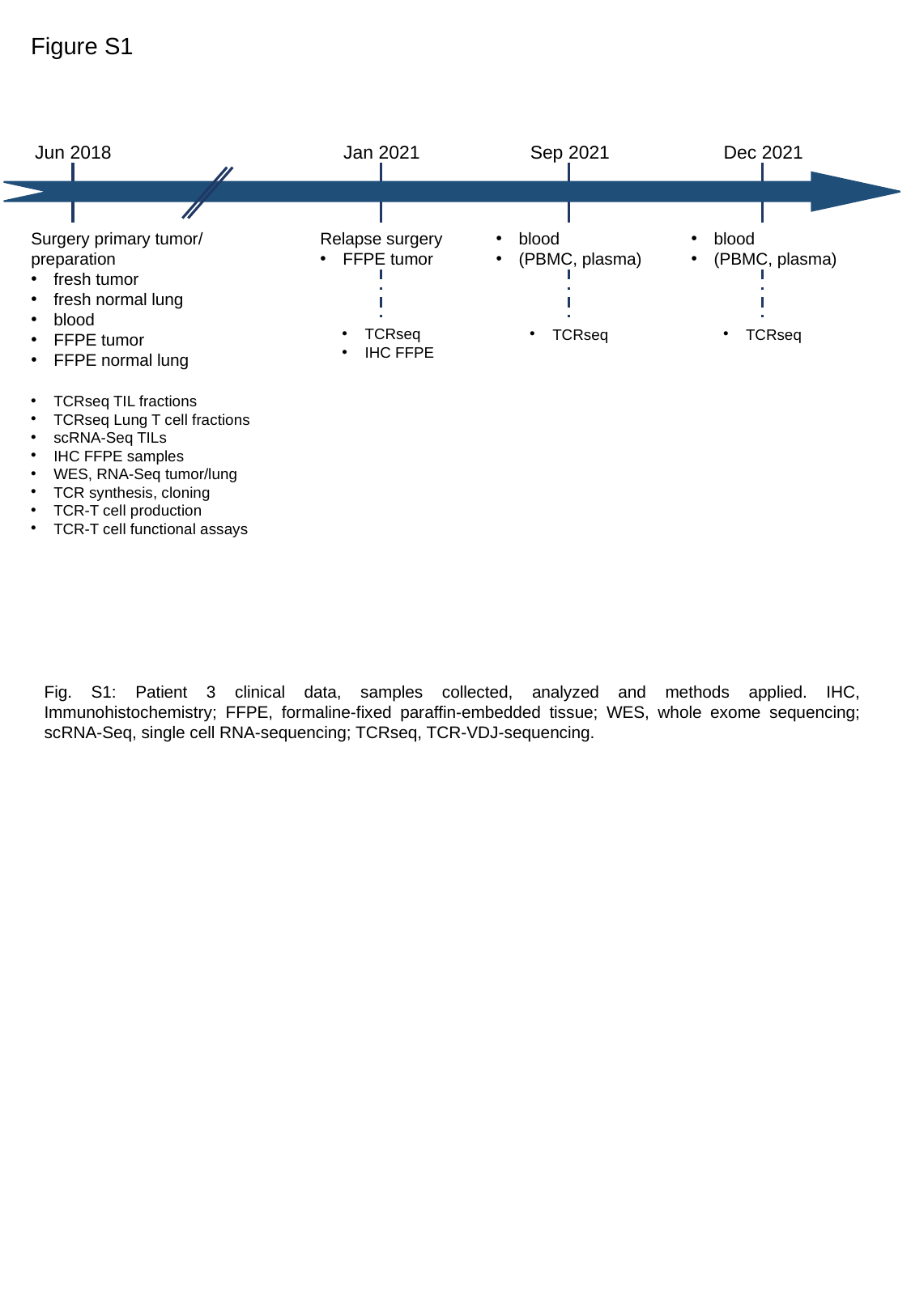

Figure S1
Jun 2018
Jan 2021
Sep 2021
Dec 2021
Surgery primary tumor/
preparation
fresh tumor
fresh normal lung
blood
FFPE tumor
FFPE normal lung
Relapse surgery
FFPE tumor
blood
(PBMC, plasma)
blood
(PBMC, plasma)
TCRseq
IHC FFPE
TCRseq
TCRseq
TCRseq TIL fractions
TCRseq Lung T cell fractions
scRNA-Seq TILs
IHC FFPE samples
WES, RNA-Seq tumor/lung
TCR synthesis, cloning
TCR-T cell production
TCR-T cell functional assays
Fig. S1: Patient 3 clinical data, samples collected, analyzed and methods applied. IHC, Immunohistochemistry; FFPE, formaline-fixed paraffin-embedded tissue; WES, whole exome sequencing; scRNA-Seq, single cell RNA-sequencing; TCRseq, TCR-VDJ-sequencing.

### Slide 3
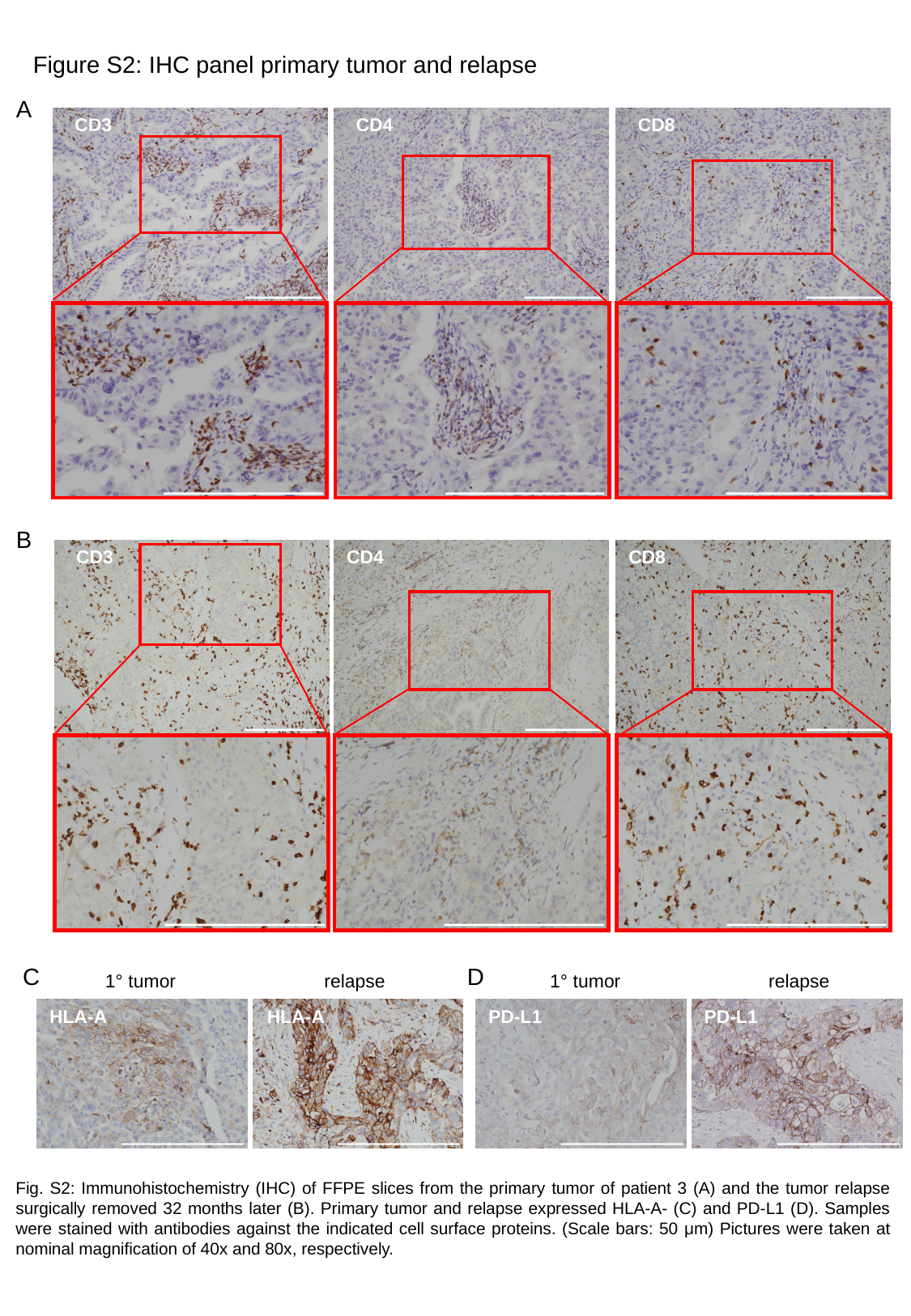

Figure S2: IHC panel primary tumor and relapse
A
CD3
CD4
CD8
B
CD3
CD4
CD8
 C
 D
1° tumor
HLA-A
relapse
HLA-A
1° tumor
PD-L1
relapse
PD-L1
Fig. S2: Immunohistochemistry (IHC) of FFPE slices from the primary tumor of patient 3 (A) and the tumor relapse surgically removed 32 months later (B). Primary tumor and relapse expressed HLA-A- (C) and PD-L1 (D). Samples were stained with antibodies against the indicated cell surface proteins. (Scale bars: 50 μm) Pictures were taken at nominal magnification of 40x and 80x, respectively.

### Slide 4
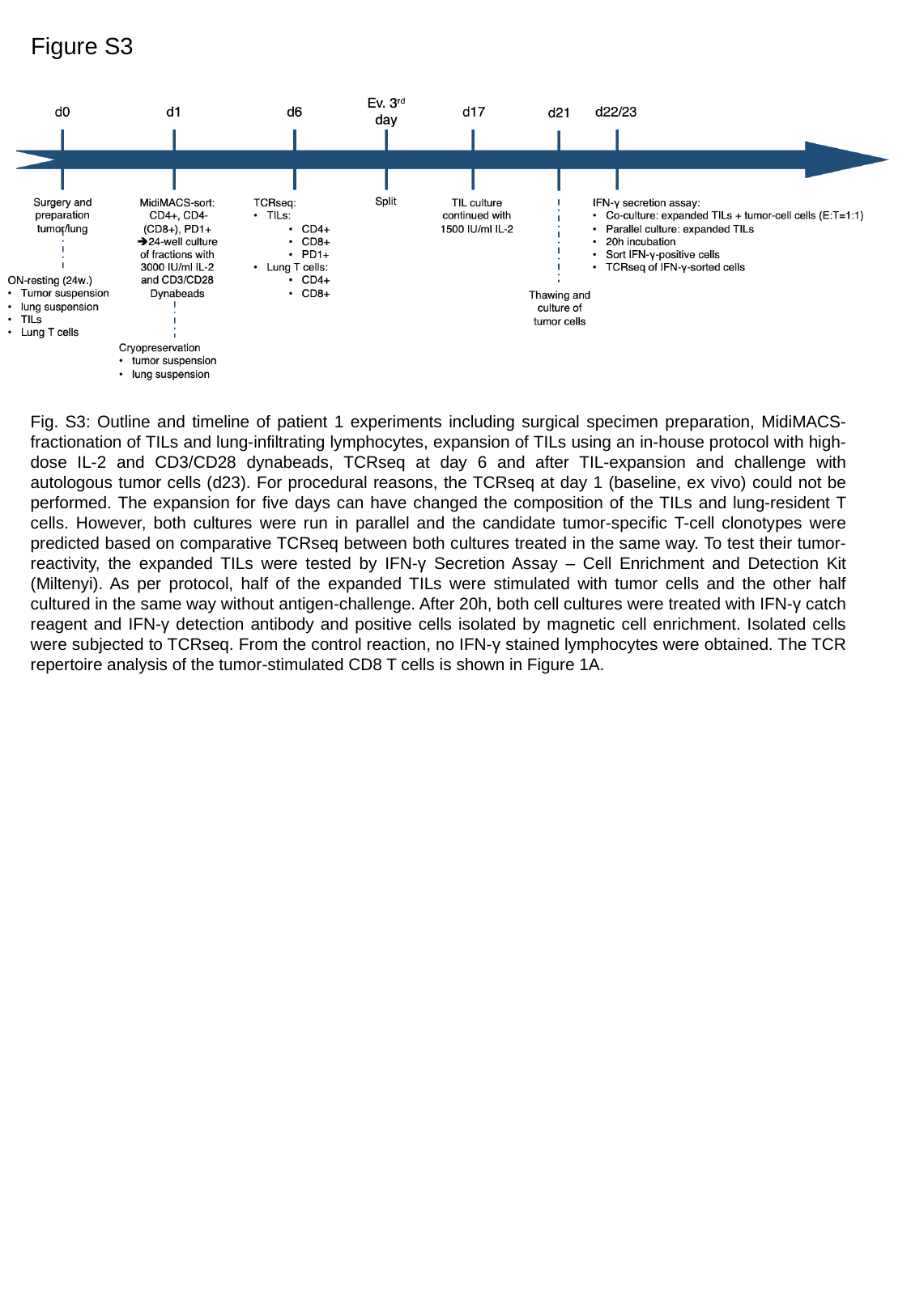

Figure S3
Fig. S3: Outline and timeline of patient 1 experiments including surgical specimen preparation, MidiMACS-fractionation of TILs and lung-infiltrating lymphocytes, expansion of TILs using an in-house protocol with high-dose IL-2 and CD3/CD28 dynabeads, TCRseq at day 6 and after TIL-expansion and challenge with autologous tumor cells (d23). For procedural reasons, the TCRseq at day 1 (baseline, ex vivo) could not be performed. The expansion for five days can have changed the composition of the TILs and lung-resident T cells. However, both cultures were run in parallel and the candidate tumor-specific T-cell clonotypes were predicted based on comparative TCRseq between both cultures treated in the same way. To test their tumor-reactivity, the expanded TILs were tested by IFN-γ Secretion Assay – Cell Enrichment and Detection Kit (Miltenyi). As per protocol, half of the expanded TILs were stimulated with tumor cells and the other half cultured in the same way without antigen-challenge. After 20h, both cell cultures were treated with IFN-γ catch reagent and IFN-γ detection antibody and positive cells isolated by magnetic cell enrichment. Isolated cells were subjected to TCRseq. From the control reaction, no IFN-γ stained lymphocytes were obtained. The TCR repertoire analysis of the tumor-stimulated CD8 T cells is shown in Figure 1A.

### Slide 5
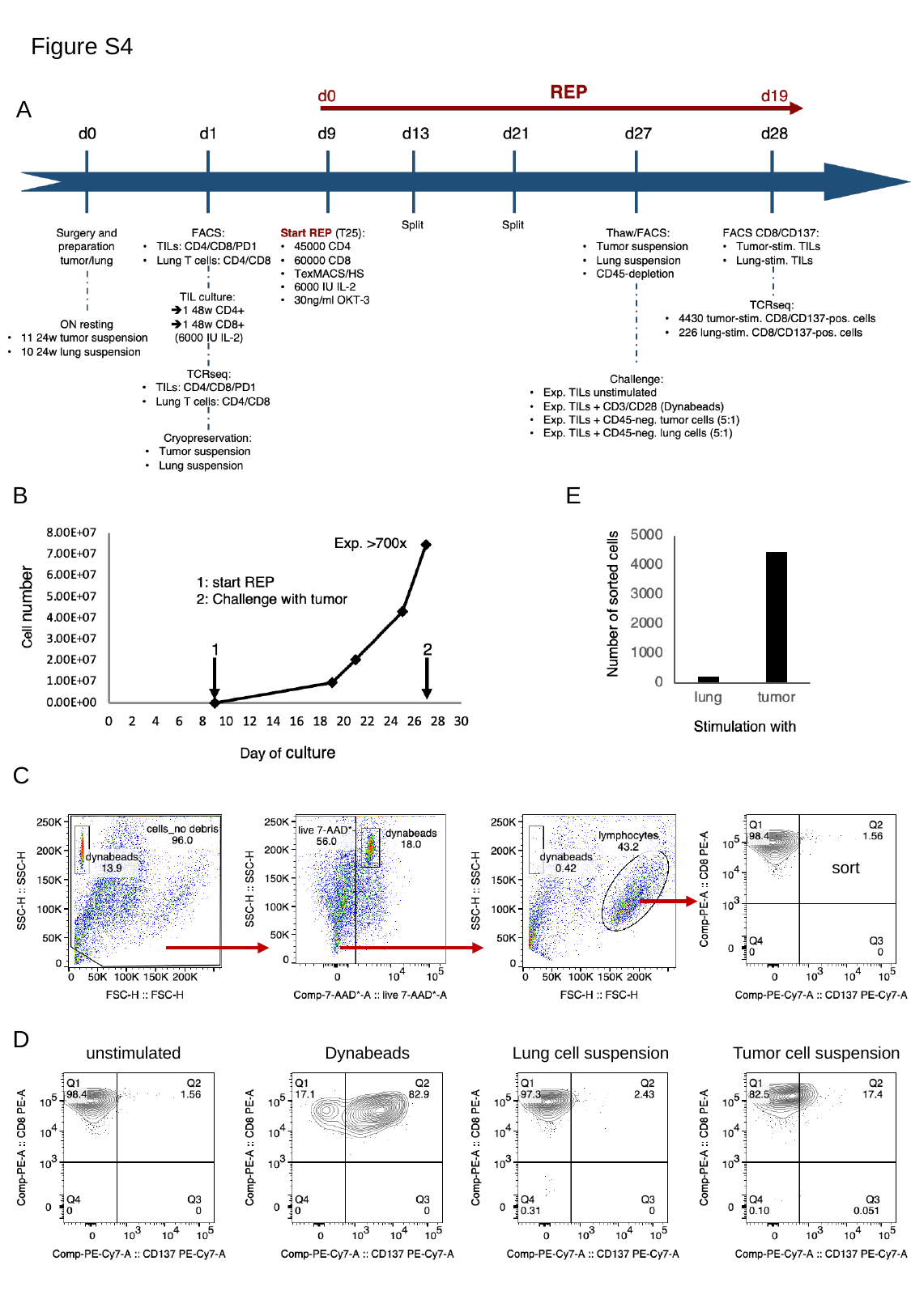

Figure S4
A
B
E
C
sort
D
unstimulated
Dynabeads
Lung cell suspension
Tumor cell suspension

### Slide 6
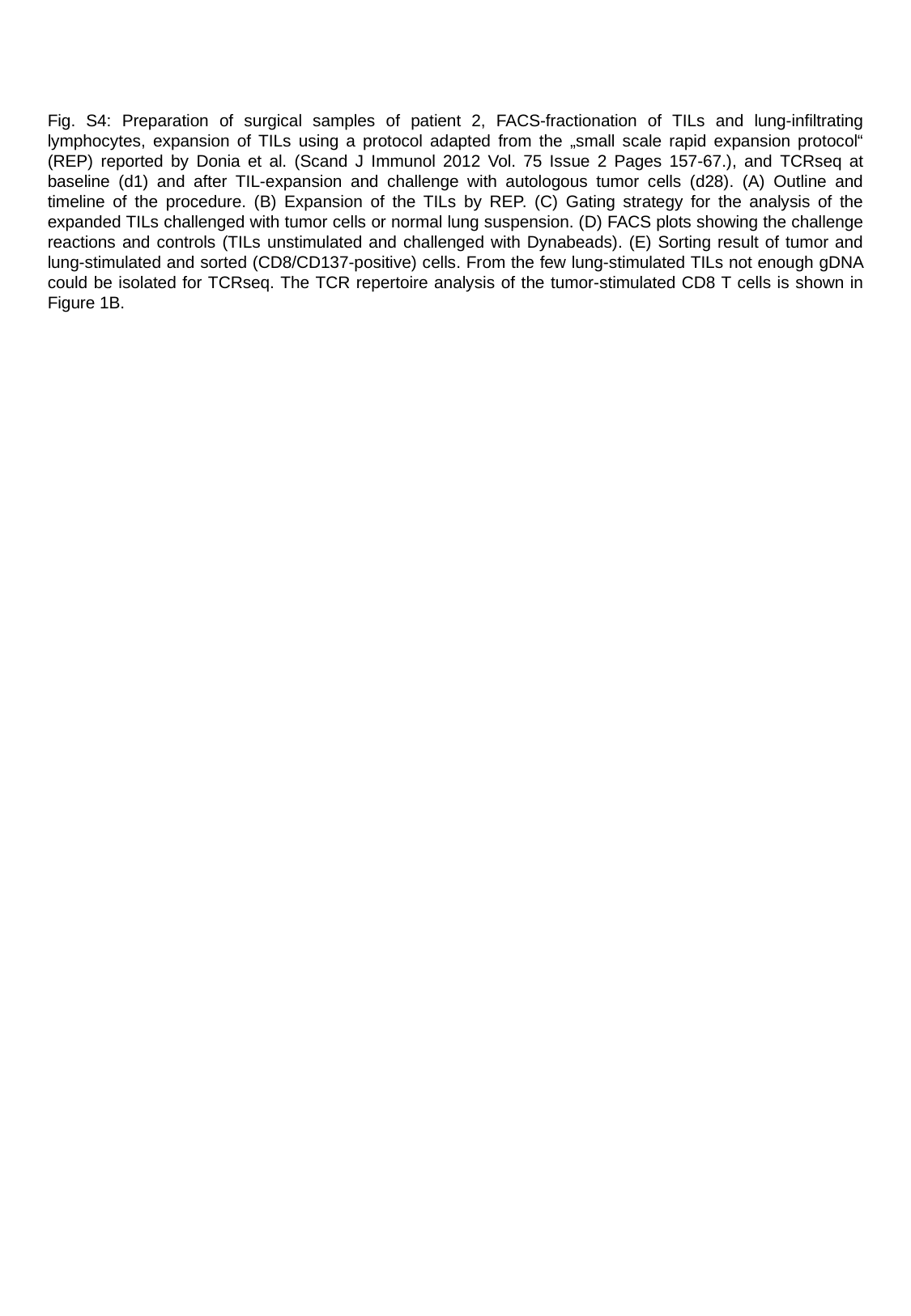

Fig. S4: Preparation of surgical samples of patient 2, FACS-fractionation of TILs and lung-infiltrating lymphocytes, expansion of TILs using a protocol adapted from the „small scale rapid expansion protocol“ (REP) reported by Donia et al. (Scand J Immunol 2012 Vol. 75 Issue 2 Pages 157-67.), and TCRseq at baseline (d1) and after TIL-expansion and challenge with autologous tumor cells (d28). (A) Outline and timeline of the procedure. (B) Expansion of the TILs by REP. (C) Gating strategy for the analysis of the expanded TILs challenged with tumor cells or normal lung suspension. (D) FACS plots showing the challenge reactions and controls (TILs unstimulated and challenged with Dynabeads). (E) Sorting result of tumor and lung-stimulated and sorted (CD8/CD137-positive) cells. From the few lung-stimulated TILs not enough gDNA could be isolated for TCRseq. The TCR repertoire analysis of the tumor-stimulated CD8 T cells is shown in Figure 1B.

### Slide 7
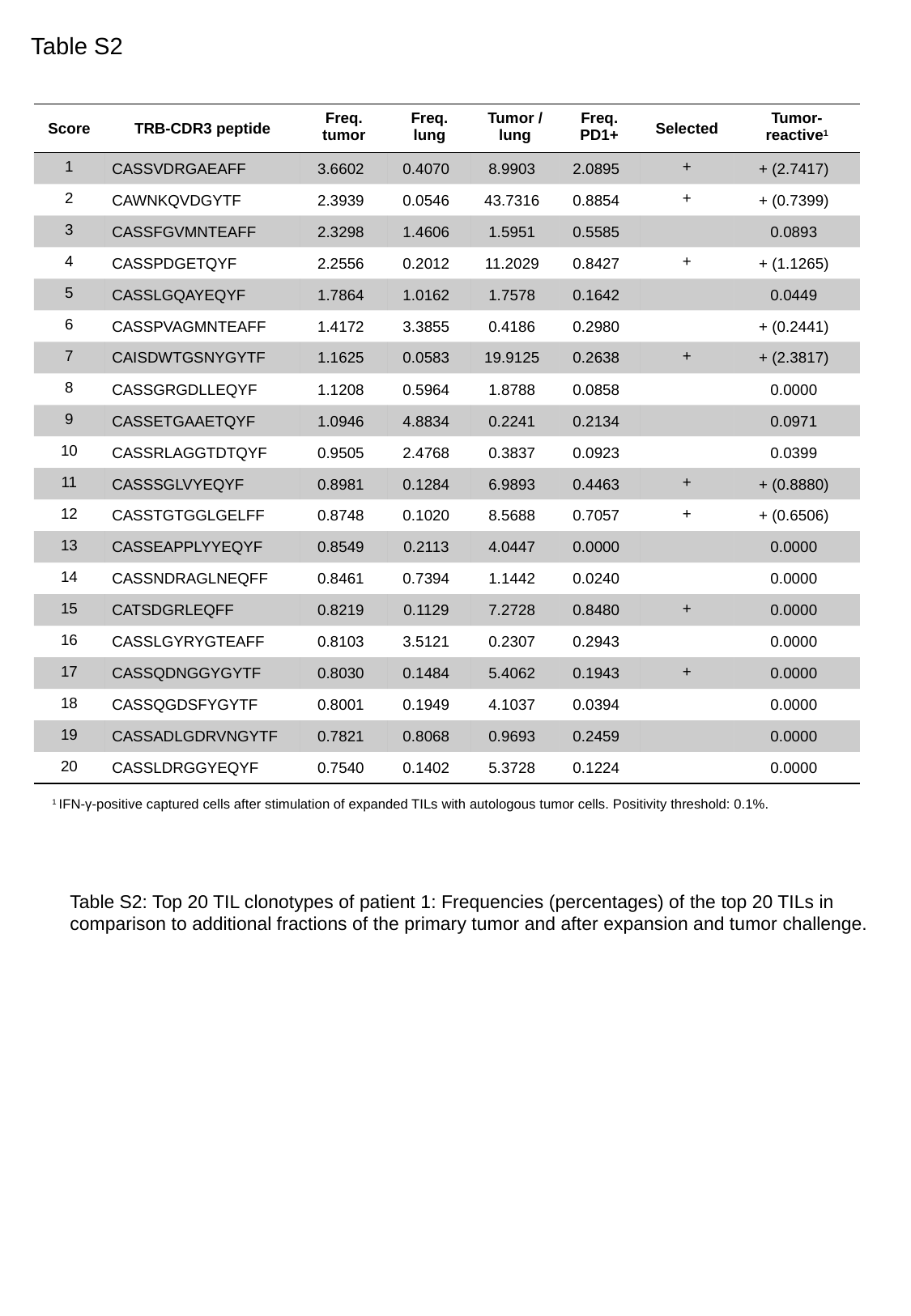

Table S2
| Score | TRB-CDR3 peptide | Freq. tumor | Freq. lung | Tumor / lung | Freq. PD1+ | Selected | Tumor-reactive^1^ |
| --- | --- | --- | --- | --- | --- | --- | --- |
| 1 | CASSVDRGAEAFF | 3.6602 | 0.4070 | 8.9903 | 2.0895 | + | + (2.7417) |
| 2 | CAWNKQVDGYTF | 2.3939 | 0.0546 | 43.7316 | 0.8854 | + | + (0.7399) |
| 3 | CASSFGVMNTEAFF | 2.3298 | 1.4606 | 1.5951 | 0.5585 | | 0.0893 |
| 4 | CASSPDGETQYF | 2.2556 | 0.2012 | 11.2029 | 0.8427 | + | + (1.1265) |
| 5 | CASSLGQAYEQYF | 1.7864 | 1.0162 | 1.7578 | 0.1642 | | 0.0449 |
| 6 | CASSPVAGMNTEAFF | 1.4172 | 3.3855 | 0.4186 | 0.2980 | | + (0.2441) |
| 7 | CAISDWTGSNYGYTF | 1.1625 | 0.0583 | 19.9125 | 0.2638 | + | + (2.3817) |
| 8 | CASSGRGDLLEQYF | 1.1208 | 0.5964 | 1.8788 | 0.0858 | | 0.0000 |
| 9 | CASSETGAAETQYF | 1.0946 | 4.8834 | 0.2241 | 0.2134 | | 0.0971 |
| 10 | CASSRLAGGTDTQYF | 0.9505 | 2.4768 | 0.3837 | 0.0923 | | 0.0399 |
| 11 | CASSSGLVYEQYF | 0.8981 | 0.1284 | 6.9893 | 0.4463 | + | + (0.8880) |
| 12 | CASSTGTGGLGELFF | 0.8748 | 0.1020 | 8.5688 | 0.7057 | + | + (0.6506) |
| 13 | CASSEAPPLYYEQYF | 0.8549 | 0.2113 | 4.0447 | 0.0000 | | 0.0000 |
| 14 | CASSNDRAGLNEQFF | 0.8461 | 0.7394 | 1.1442 | 0.0240 | | 0.0000 |
| 15 | CATSDGRLEQFF | 0.8219 | 0.1129 | 7.2728 | 0.8480 | + | 0.0000 |
| 16 | CASSLGYRYGTEAFF | 0.8103 | 3.5121 | 0.2307 | 0.2943 | | 0.0000 |
| 17 | CASSQDNGGYGYTF | 0.8030 | 0.1484 | 5.4062 | 0.1943 | + | 0.0000 |
| 18 | CASSQGDSFYGYTF | 0.8001 | 0.1949 | 4.1037 | 0.0394 | | 0.0000 |
| 19 | CASSADLGDRVNGYTF | 0.7821 | 0.8068 | 0.9693 | 0.2459 | | 0.0000 |
| 20 | CASSLDRGGYEQYF | 0.7540 | 0.1402 | 5.3728 | 0.1224 | | 0.0000 |
^1^ IFN-γ-positive captured cells after stimulation of expanded TILs with autologous tumor cells. Positivity threshold: 0.1%.
Table S2: Top 20 TIL clonotypes of patient 1: Frequencies (percentages) of the top 20 TILs in comparison to additional fractions of the primary tumor and after expansion and tumor challenge.

### Slide 8
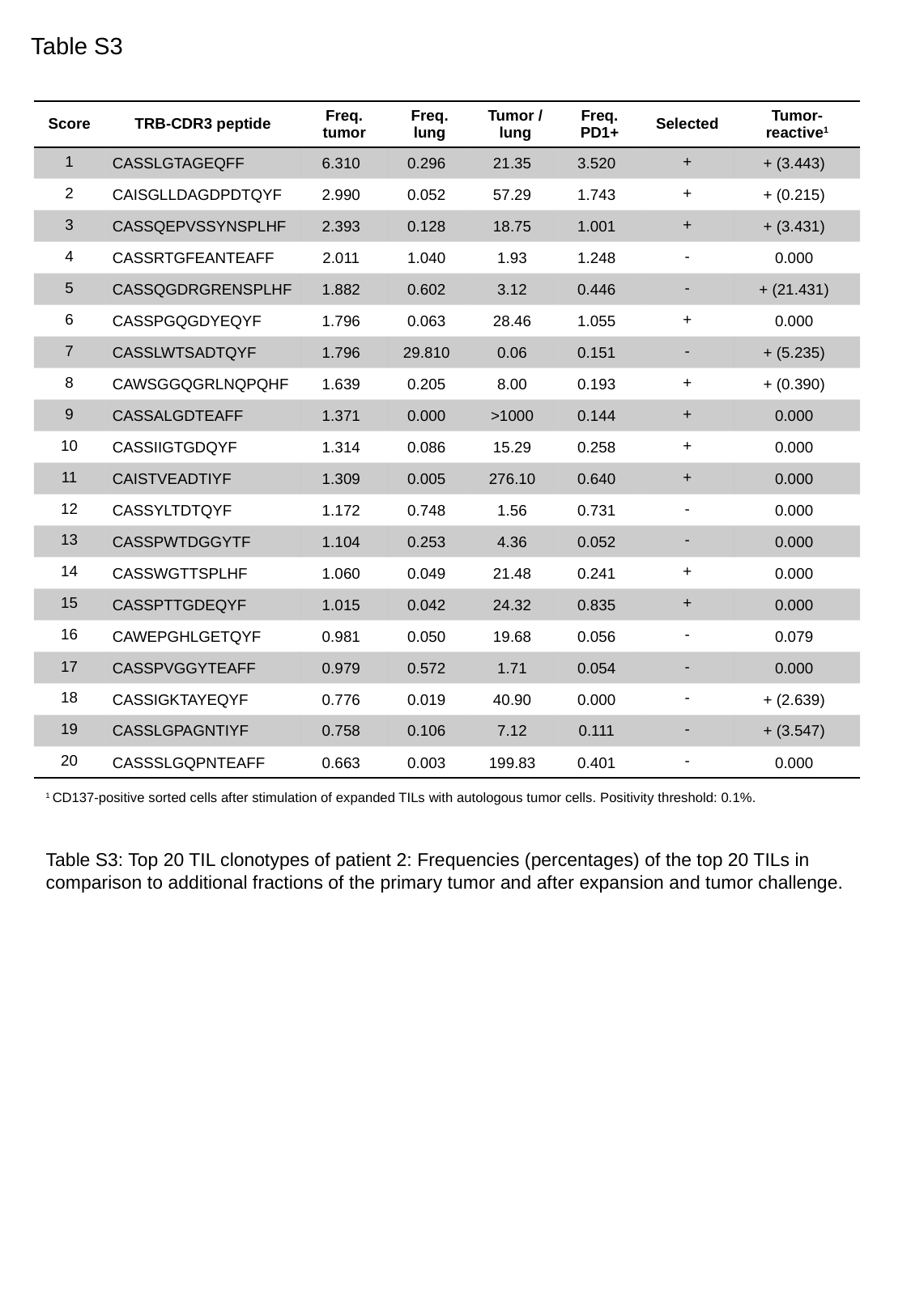

Table S3
| Score | TRB-CDR3 peptide | Freq. tumor | Freq. lung | Tumor / lung | Freq. PD1+ | Selected | Tumor-reactive^1^ |
| --- | --- | --- | --- | --- | --- | --- | --- |
| 1 | CASSLGTAGEQFF | 6.310 | 0.296 | 21.35 | 3.520 | + | + (3.443) |
| 2 | CAISGLLDAGDPDTQYF | 2.990 | 0.052 | 57.29 | 1.743 | + | + (0.215) |
| 3 | CASSQEPVSSYNSPLHF | 2.393 | 0.128 | 18.75 | 1.001 | + | + (3.431) |
| 4 | CASSRTGFEANTEAFF | 2.011 | 1.040 | 1.93 | 1.248 | - | 0.000 |
| 5 | CASSQGDRGRENSPLHF | 1.882 | 0.602 | 3.12 | 0.446 | - | + (21.431) |
| 6 | CASSPGQGDYEQYF | 1.796 | 0.063 | 28.46 | 1.055 | + | 0.000 |
| 7 | CASSLWTSADTQYF | 1.796 | 29.810 | 0.06 | 0.151 | - | + (5.235) |
| 8 | CAWSGGQGRLNQPQHF | 1.639 | 0.205 | 8.00 | 0.193 | + | + (0.390) |
| 9 | CASSALGDTEAFF | 1.371 | 0.000 | >1000 | 0.144 | + | 0.000 |
| 10 | CASSIIGTGDQYF | 1.314 | 0.086 | 15.29 | 0.258 | + | 0.000 |
| 11 | CAISTVEADTIYF | 1.309 | 0.005 | 276.10 | 0.640 | + | 0.000 |
| 12 | CASSYLTDTQYF | 1.172 | 0.748 | 1.56 | 0.731 | - | 0.000 |
| 13 | CASSPWTDGGYTF | 1.104 | 0.253 | 4.36 | 0.052 | - | 0.000 |
| 14 | CASSWGTTSPLHF | 1.060 | 0.049 | 21.48 | 0.241 | + | 0.000 |
| 15 | CASSPTTGDEQYF | 1.015 | 0.042 | 24.32 | 0.835 | + | 0.000 |
| 16 | CAWEPGHLGETQYF | 0.981 | 0.050 | 19.68 | 0.056 | - | 0.079 |
| 17 | CASSPVGGYTEAFF | 0.979 | 0.572 | 1.71 | 0.054 | - | 0.000 |
| 18 | CASSIGKTAYEQYF | 0.776 | 0.019 | 40.90 | 0.000 | - | + (2.639) |
| 19 | CASSLGPAGNTIYF | 0.758 | 0.106 | 7.12 | 0.111 | - | + (3.547) |
| 20 | CASSSLGQPNTEAFF | 0.663 | 0.003 | 199.83 | 0.401 | - | 0.000 |
^1^ CD137-positive sorted cells after stimulation of expanded TILs with autologous tumor cells. Positivity threshold: 0.1%.
Table S3: Top 20 TIL clonotypes of patient 2: Frequencies (percentages) of the top 20 TILs in comparison to additional fractions of the primary tumor and after expansion and tumor challenge.

### Slide 9
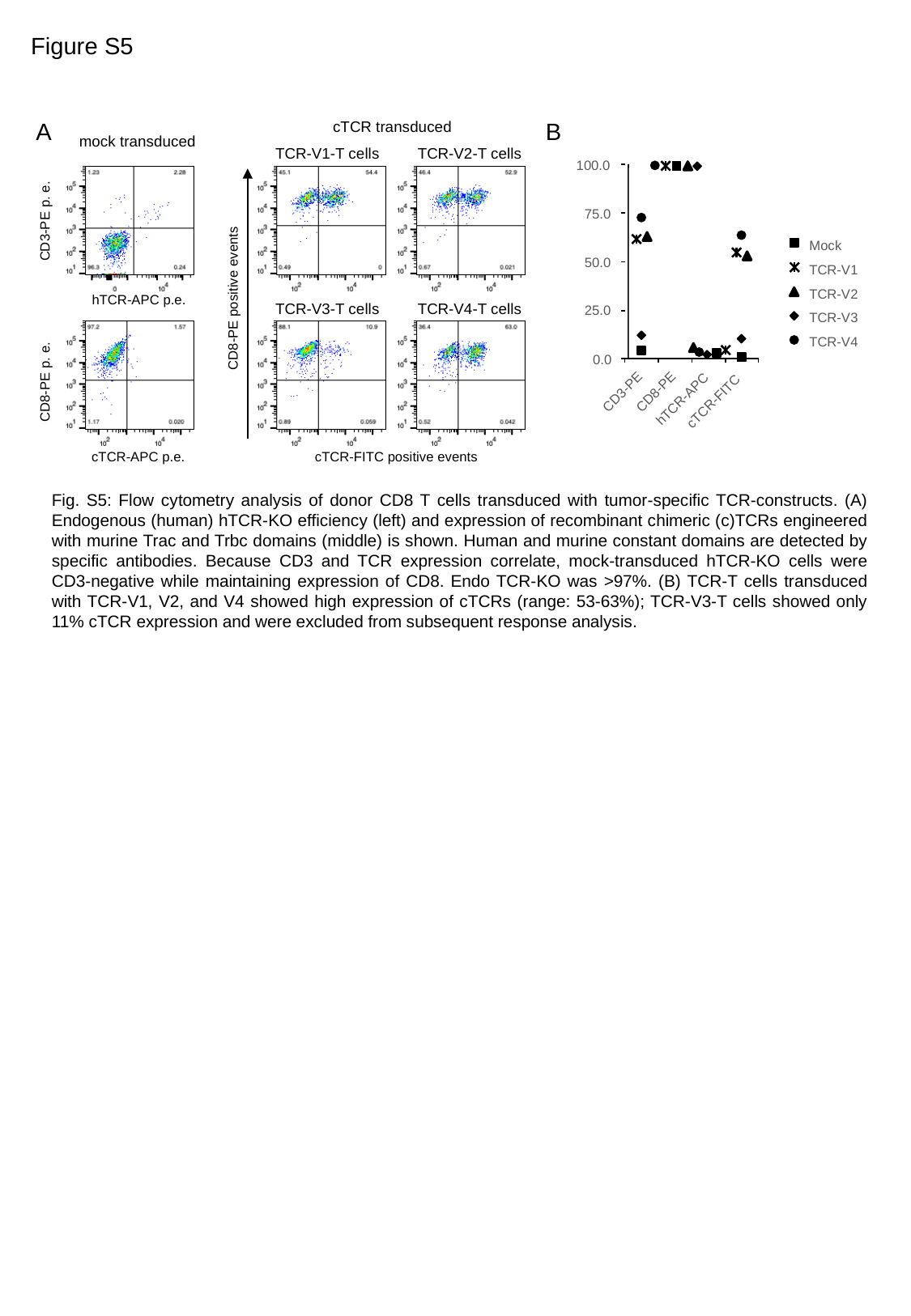

Figure S5
A
B
cTCR transduced
TCR-V1-T cells
TCR-V2-T cells
CD8-PE positive events
TCR-V3-T cells
TCR-V4-T cells
cTCR-FITC positive events
mock transduced
CD3-PE p. e.
hTCR-APC p.e.
CD8-PE p. e.
cTCR-APC p.e.
100.0
75.0
50.0
25.0
0.0
CD3-PE
CD8-PE
hTCR-APC
cTCR-FITC
Mock
TCR-V1
TCR-V2
TCR-V3
TCR-V4
Fig. S5: Flow cytometry analysis of donor CD8 T cells transduced with tumor-specific TCR-constructs. (A) Endogenous (human) hTCR-KO efficiency (left) and expression of recombinant chimeric (c)TCRs engineered with murine Trac and Trbc domains (middle) is shown. Human and murine constant domains are detected by specific antibodies. Because CD3 and TCR expression correlate, mock-transduced hTCR-KO cells were CD3-negative while maintaining expression of CD8. Endo TCR-KO was >97%. (B) TCR-T cells transduced with TCR-V1, V2, and V4 showed high expression of cTCRs (range: 53-63%); TCR-V3-T cells showed only 11% cTCR expression and were excluded from subsequent response analysis.

### Slide 10
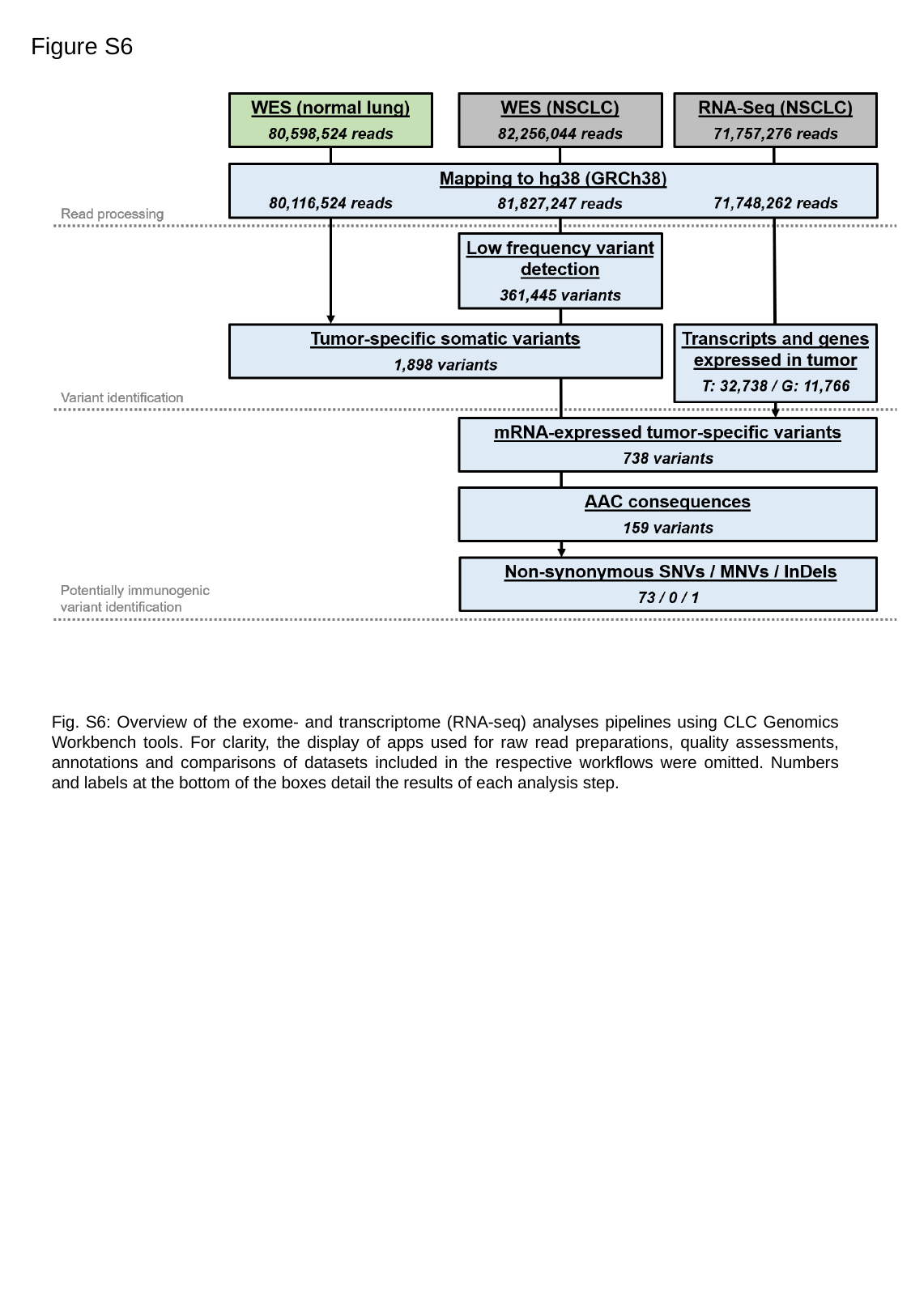

Figure S6
Fig. S6: Overview of the exome- and transcriptome (RNA-seq) analyses pipelines using CLC Genomics Workbench tools. For clarity, the display of apps used for raw read preparations, quality assessments, annotations and comparisons of datasets included in the respective workflows were omitted. Numbers and labels at the bottom of the boxes detail the results of each analysis step.

### Slide 11
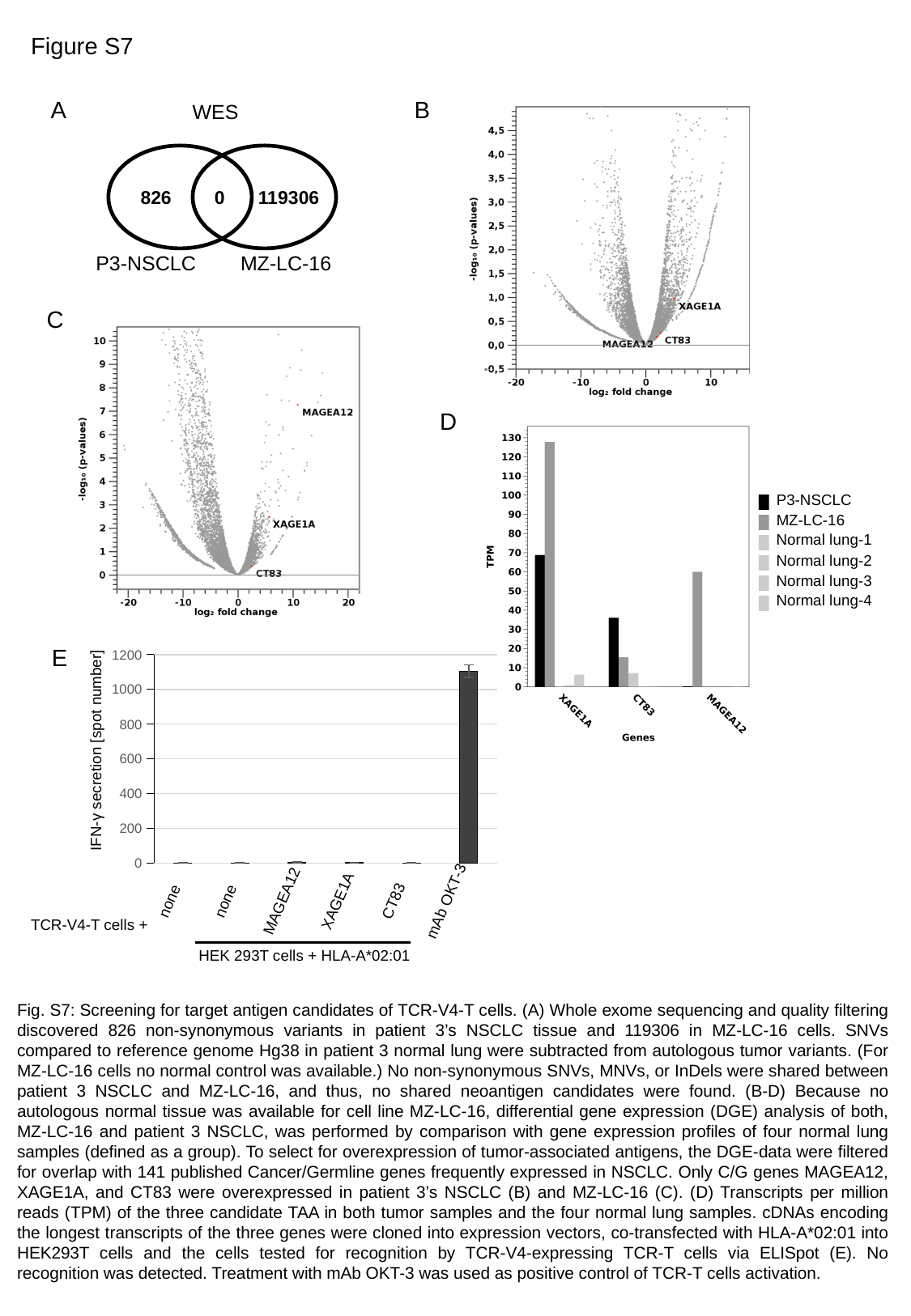

Figure S7
A
WES
826
0
119306
P3-NSCLC
MZ-LC-16
B
C
D
P3-NSCLC
MZ-LC-16
Normal lung-1
Normal lung-2
Normal lung-3
Normal lung-4
#### Chart
| Category | |
|---|---|
| TCR-T only | 1.0 |
| + 293T/HLA-A | 1.0 |
| + MAGE-A12 | 3.0 |
| +XAGE-A1 | 3.0 |
| +CT83 | 2.0 |
| +OKT-3 | 1105.0 |IFN-γ secretion [spot number]
none
none
MAGEA12
XAGE1A
CT83
mAb OKT-3
TCR-V4-T cells +
HEK 293T cells + HLA-A*02:01
E
Fig. S7: Screening for target antigen candidates of TCR-V4-T cells. (A) Whole exome sequencing and quality filtering discovered 826 non-synonymous variants in patient 3’s NSCLC tissue and 119306 in MZ-LC-16 cells. SNVs compared to reference genome Hg38 in patient 3 normal lung were subtracted from autologous tumor variants. (For MZ-LC-16 cells no normal control was available.) No non-synonymous SNVs, MNVs, or InDels were shared between patient 3 NSCLC and MZ-LC-16, and thus, no shared neoantigen candidates were found. (B-D) Because no autologous normal tissue was available for cell line MZ-LC-16, differential gene expression (DGE) analysis of both, MZ-LC-16 and patient 3 NSCLC, was performed by comparison with gene expression profiles of four normal lung samples (defined as a group). To select for overexpression of tumor-associated antigens, the DGE-data were filtered for overlap with 141 published Cancer/Germline genes frequently expressed in NSCLC. Only C/G genes MAGEA12, XAGE1A, and CT83 were overexpressed in patient 3’s NSCLC (B) and MZ-LC-16 (C). (D) Transcripts per million reads (TPM) of the three candidate TAA in both tumor samples and the four normal lung samples. cDNAs encoding the longest transcripts of the three genes were cloned into expression vectors, co-transfected with HLA-A*02:01 into HEK293T cells and the cells tested for recognition by TCR-V4-expressing TCR-T cells via ELISpot (E). No recognition was detected. Treatment with mAb OKT-3 was used as positive control of TCR-T cells activation.

### Slide 12
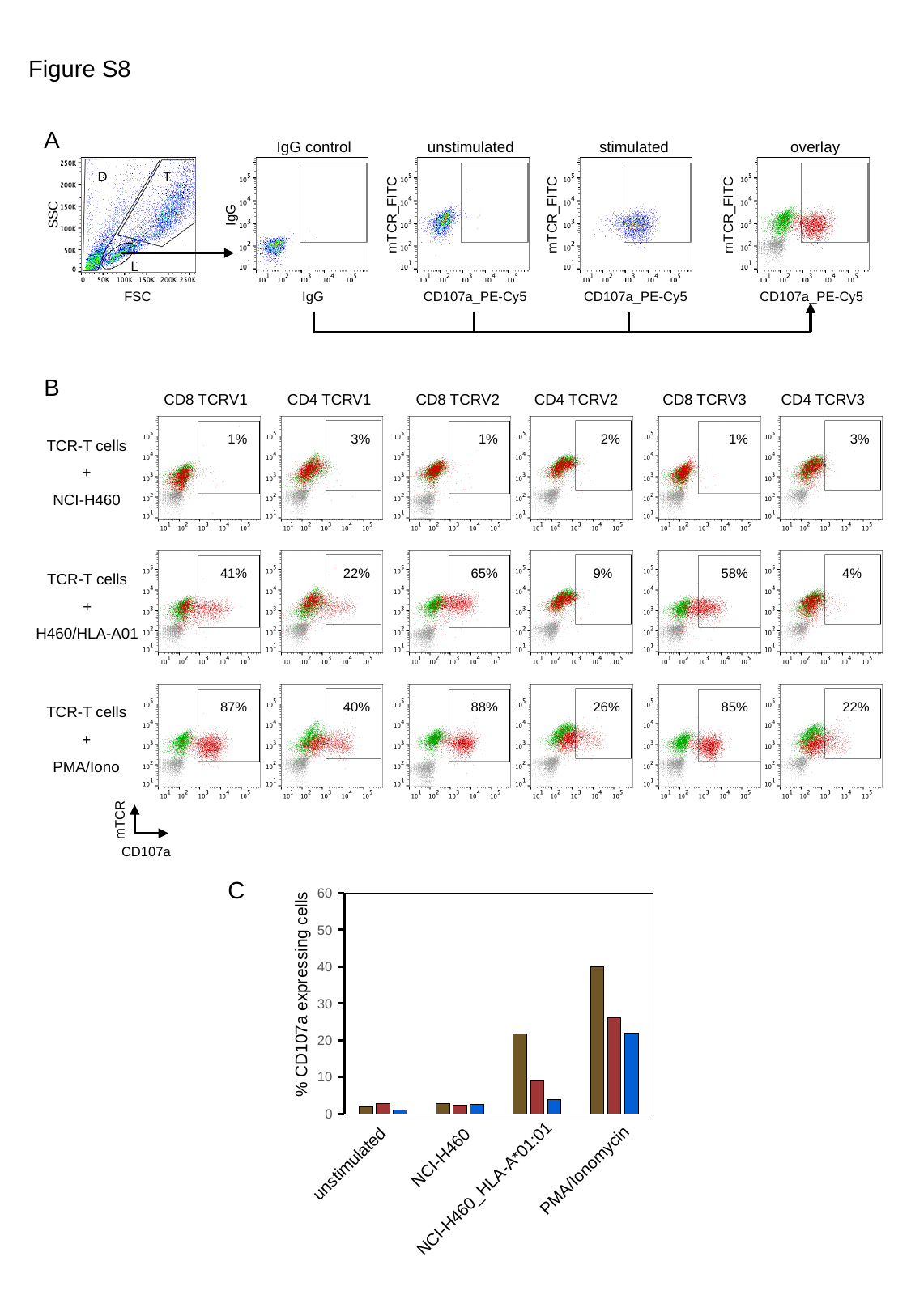

Figure S8
A
IgG control
unstimulated
stimulated
overlay
D
T
SSC
IgG
mTCR_FITC
mTCR_FITC
mTCR_FITC
L
FSC
IgG
CD107a_PE-Cy5
CD107a_PE-Cy5
CD107a_PE-Cy5
B
CD8 TCRV1
CD4 TCRV1
CD8 TCRV2
CD4 TCRV2
CD8 TCRV3
CD4 TCRV3
1%
41%
87%
3%
22%
40%
1%
65%
88%
2%
9%
26%
1%
58%
85%
3%
4%
22%
TCR-T cells
+
NCI-H460
TCR-T cells
+
H460/HLA-A01
TCR-T cells
+
PMA/Iono
C
#### Chart
| Category | TCR-V1 T | TCR-V2 T | TCR-V3 T |
|---|---|---|---|
| unstimulated | 1.94 | 2.81 | 1.12 |
| NCI-H460 | 2.83 | 2.45 | 2.62 |
| NCI-H460_HLA-A01 | 21.7 | 9.0 | 3.95 |
| PMA/Ionomycin | 40.0 | 26.0 | 22.0 |% CD107a expressing cells
NCI-H460
unstimulated
PMA/Ionomycin
NCI-H460_HLA-A*01:01
mTCR
CD107a

### Slide 13
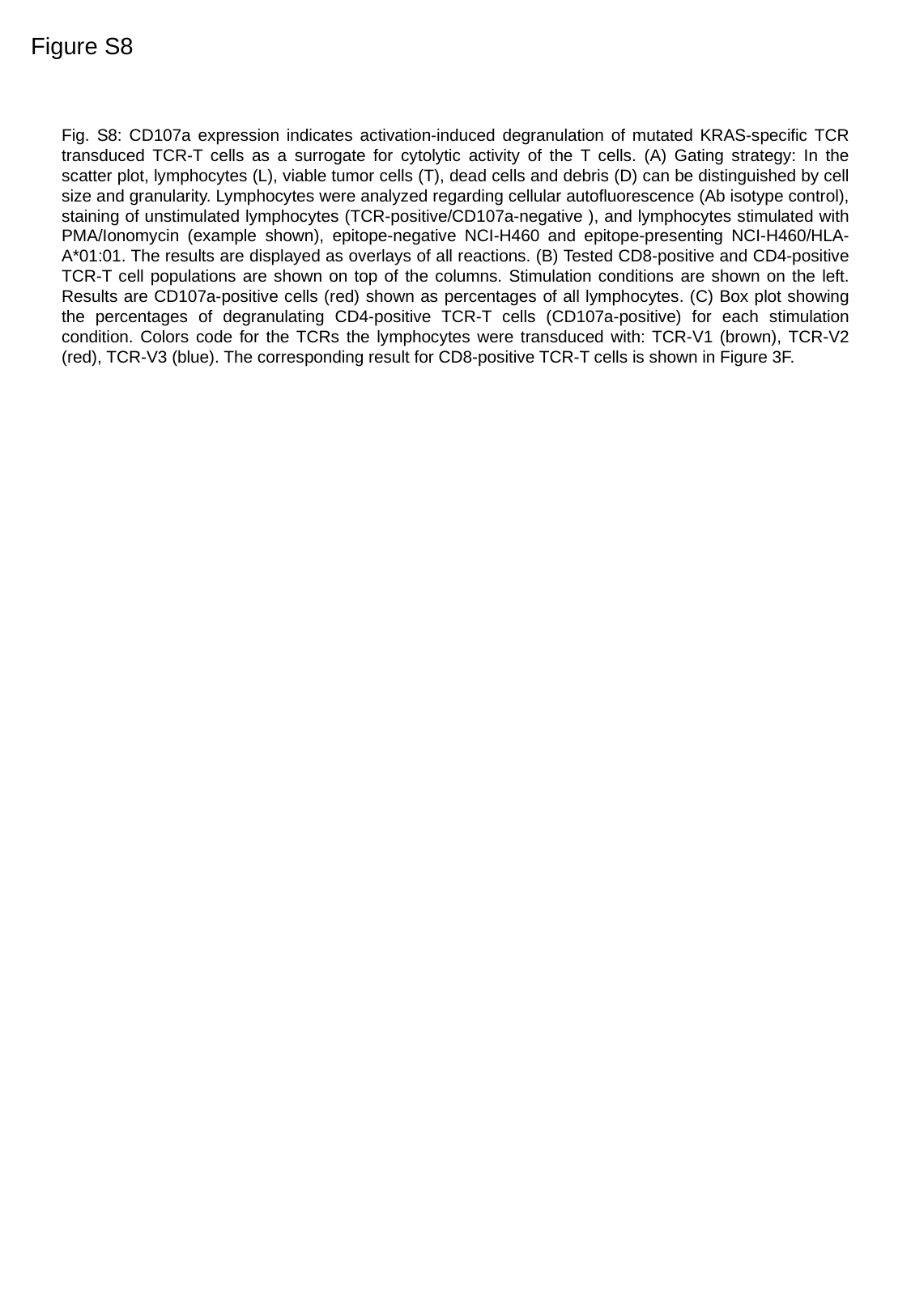

Figure S8
Fig. S8: CD107a expression indicates activation-induced degranulation of mutated KRAS-specific TCR transduced TCR-T cells as a surrogate for cytolytic activity of the T cells. (A) Gating strategy: In the scatter plot, lymphocytes (L), viable tumor cells (T), dead cells and debris (D) can be distinguished by cell size and granularity. Lymphocytes were analyzed regarding cellular autofluorescence (Ab isotype control), staining of unstimulated lymphocytes (TCR-positive/CD107a-negative ), and lymphocytes stimulated with PMA/Ionomycin (example shown), epitope-negative NCI-H460 and epitope-presenting NCI-H460/HLA-A*01:01. The results are displayed as overlays of all reactions. (B) Tested CD8-positive and CD4-positive TCR-T cell populations are shown on top of the columns. Stimulation conditions are shown on the left. Results are CD107a-positive cells (red) shown as percentages of all lymphocytes. (C) Box plot showing the percentages of degranulating CD4-positive TCR-T cells (CD107a-positive) for each stimulation condition. Colors code for the TCRs the lymphocytes were transduced with: TCR-V1 (brown), TCR-V2 (red), TCR-V3 (blue). The corresponding result for CD8-positive TCR-T cells is shown in Figure 3F.

### Slide 14
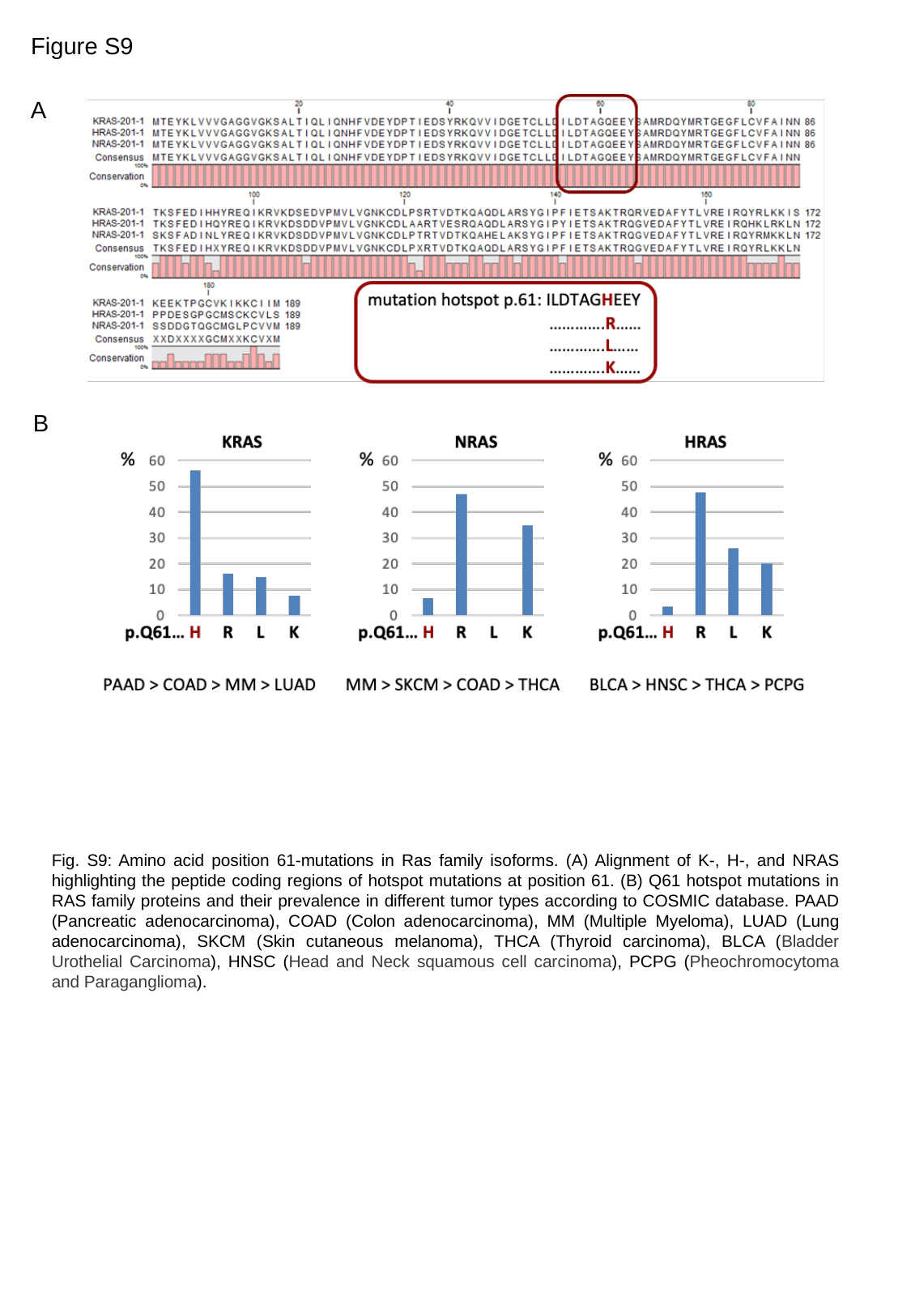

Figure S9
A
B
Fig. S9: Amino acid position 61-mutations in Ras family isoforms. (A) Alignment of K-, H-, and NRAS highlighting the peptide coding regions of hotspot mutations at position 61. (B) Q61 hotspot mutations in RAS family proteins and their prevalence in different tumor types according to COSMIC database. PAAD (Pancreatic adenocarcinoma), COAD (Colon adenocarcinoma), MM (Multiple Myeloma), LUAD (Lung adenocarcinoma), SKCM (Skin cutaneous melanoma), THCA (Thyroid carcinoma), BLCA (Bladder Urothelial Carcinoma), HNSC (Head and Neck squamous cell carcinoma), PCPG (Pheochromocytoma and Paraganglioma).

### Slide 15
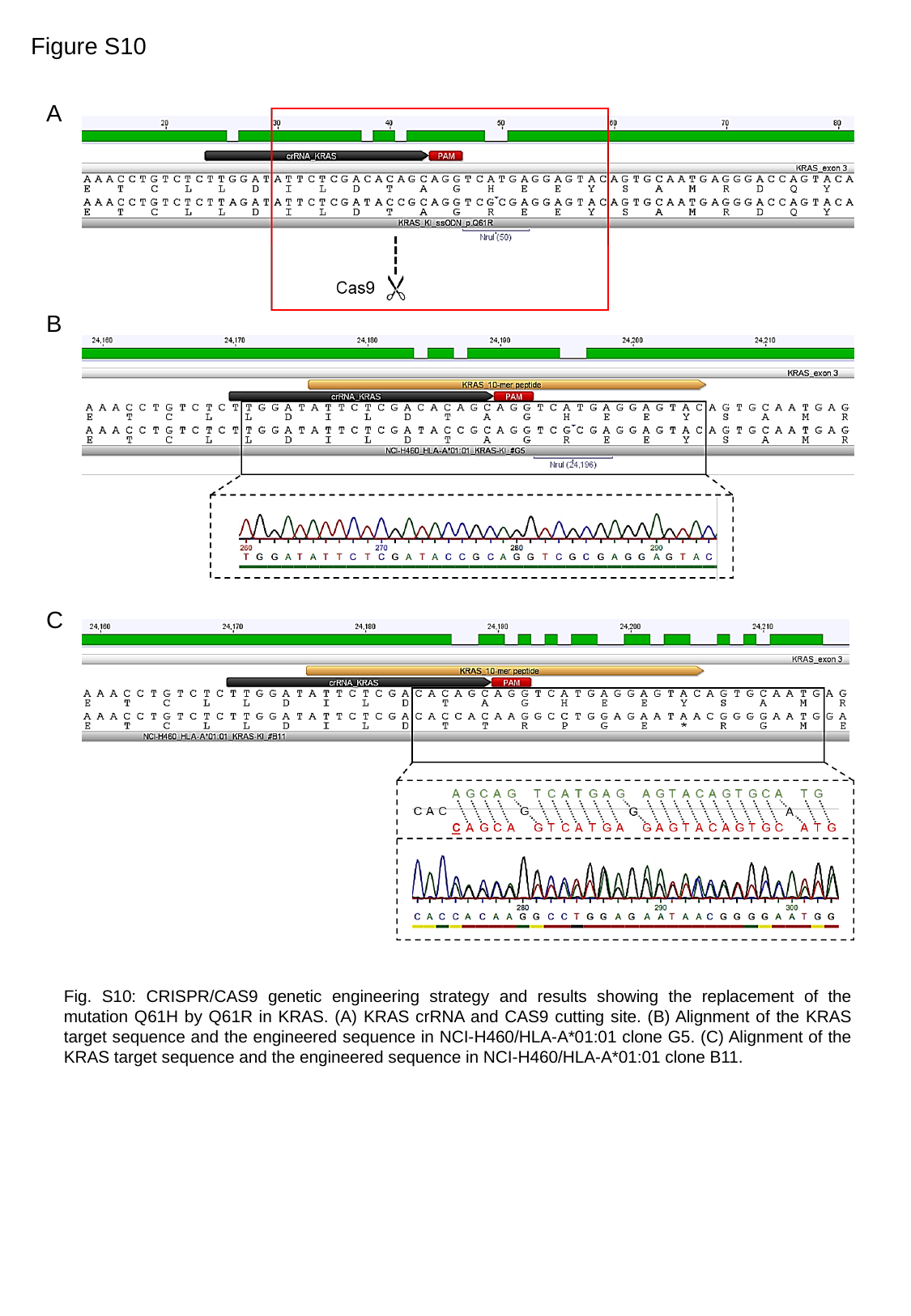

Figure S10
A
B
C
Fig. S10: CRISPR/CAS9 genetic engineering strategy and results showing the replacement of the mutation Q61H by Q61R in KRAS. (A) KRAS crRNA and CAS9 cutting site. (B) Alignment of the KRAS target sequence and the engineered sequence in NCI-H460/HLA-A*01:01 clone G5. (C) Alignment of the KRAS target sequence and the engineered sequence in NCI-H460/HLA-A*01:01 clone B11.
